# The nasopharyngeal, ruminal, and vaginal microbiota and the core taxa shared across these microbiomes in virgin yearling heifers exposed to divergent in utero nutrition during their first trimester of gestation and in pregnant beef heifers in response to mineral supplementation

**DOI:** 10.1101/2021.06.03.446997

**Authors:** Samat Amat, Devin B. Holman, Kaycie Schmidt, Ana Clara B. Menezes, Friederike Baumgaertner, Thomas Winders, James D. Kirsch, TingTing Liu, Timothy D. Schwinghamer, Kevin K. Sedivec, Carl R. Dahlen

**Affiliations:** Department of Microbiological Sciences, North Dakota State University, Fargo, ND, 58108, USA; Lacombe Research and Development Centre, Agriculture and Agri-Food Canada, 6000 C & E Trail, Lacombe, AB, T4L 1W1, Canada; Department of Animal Sciences, North Dakota State University, Fargo, ND, 58102, USA; Agriculture and Agri-Food Canada, Lethbridge Research and Development Centre, Lethbridge, AB, Canada; Central Grasslands Research Extension Center, North Dakota State University, Streeter, ND, 58483, USA

**Keywords:** Beef heifers, Core taxa, Maternal utrition, Nasopharyngeal microbiota, Offspring, Ruminal microbiota, Vaginal microbiota.

## Abstract

Emerging evidence has indicated that microbial transmission from the bovine dam to her fetus may take place before birth, and that the maternal microbiota during pregnancy modulates programming of fetal metabolic and nervous system development, highlighting the potential and extended role of the maternal microbiome in calf health and development. In the present study, we characterized the nasopharyngeal, ruminal and vaginal microbiota from two cohorts of beef heifers managed at the same location: 1) virgin yearling heifers (9 months old) born from dams received gestational diets which resulted in low (LG, n = 22) or medium (MG, n = 23) weight gain during the first 84 days of gestation; and 2) pregnant replacement heifers that received a vitamin and mineral supplement (VTM, n = 17) or not (Control, n = 15) during the first 6 months of gestation. Nasopharyngeal and vaginal swabs as well as ruminal fluid were collected from both cohorts and the microbiota of each sample was assessed using 16S rRNA gene sequencing. In addition to the comparison between treatment groups within each cohort, the similarity of the microbiota of the three sample types were evaluated, and shared taxa amongst these communities were identified. The bacterial genera present in the rumen and vagina that can influence methanogenic archaeal genera were predicted using a stepwise-selected generalized linear mixed model. No significant difference was observed in the alpha and beta diversity in any of the nasopharyngeal, ruminal and vaginal microbiota between LG and MG offspring virgin heifers, or between the control and VTM pregnant heifers (*p* > 0.05). Subtle compositional changes in the vaginal microbiota in yearling heifers, and in the nasopharyngeal and ruminal microbiota of pregnant heifers were detected in response to treatments. Forty-one archaeal and bacterial OTUs were shared by over 60% of all samples from both virgin and pregnant heifers. Two taxa within the *Methanobrevibacter* genus were identified as core taxa and this genus was more relatively abundant in pregnant heifers compared to virgin heifers. Among the 25 top genera, *Prevotella* and *Prevotella* UCG-003 (negative) and *Christensenellaceae R-7* group (positive) were predicted to have a significant effect on ruminal *Methanobrevibacter* spp. The results of this study indicate that there is little impact of divergent gestational nutrition during the first trimester on the calf microbiome at 9 months postnatal, and that VTM supplementation during pregnancy may not alter the maternal microbiome. This study provides evidence that there are several microbial taxa, including methanogenic archaea, that are shared across the respiratory, gastrointestinal, and reproductive tracts, suggesting the need for a holistic evaluation of the bovine microbiota when considering potential maternal sources for seeding calves with pioneer microbiota.

## INTRODUCTION

Host genetic selection has been a primary target for improving animal health and productivity over the last several decades and has resulted in tremendous progress in both dairy and beef cattle production systems. Recently, the microbiota colonizing different mucosal surfaces of cattle have become a new target for manipulation/engineering with great potential to improve animal health and production (Huws et al., 2018; Matthews et al., 2019). Diverse and dynamic microbial communities present in the respiratory, gastrointestinal and reproductive tracts of cattle are vital to health and performance (Galvão et al., 2019; Matthews et al., 2019; Timsit et al., 2020). Among these microbial communities, the ruminal microbiota in cattle, which is the most densely populated and involved in both nutrient metabolism and immune system development, has become the primary target for manipulation/engineering (O’Hara et al., 2020).

Recent developments including the advent of high-throughput sequencing techniques, heritable ruminal microbiota compositional changes that are associated with feed efficiency (Difford et al., 2018; Li et al., 2019) and methane emission phenotypes in cattle (Difford et al., 2018), suggest that the ruminal microbiome and host genetics can be targeted independently to improve feed efficiency and mitigate enteric methane emissions from cattle. One of the challenges associated with manipulation of the ruminal microbiome in mature animals is its resiliency that allows the microbiome to revert to the original state following the cessation of an intervention (Weimer, 2015). To overcome this challenge, early life microbial programming in young ruminants was recommended and has shown some efficacy (Yáñez-Ruiz et al., 2015; Saro et al., 2018; Belanche et al., 2020). For example, Palma-Hidalgo et al. (2021) reported that the direct inoculation of fresh ruminal fluid from adult goats to kids in early life accelerated ruminal microbial community development and improved the weaning process. Early life microbial programming is based on the current dogma that microbial colonization of the rumen starts at birth, and the developing ruminal microbiota within the first 3 weeks of life is less resilient to manipulation (Yáñez-Ruiz et al., 2015). A recent study by Guzman and colleagues (2020), however, provided sequencing and culture-based evidence indicating that the intestine of calf fetus is not sterile and colonization by so-called “pioneer” microbes may occur during gestation. This is further supported by our preliminary data which suggested that colonization of the fetal intestine by archaea and bacteria may take place within the first 12 weeks of gestation in cattle (Amat et al., unpublished data). These observations highlight the potential and extended role of the maternal microbiome in calf ruminal microbiome development.

Although the role of maternal nutrition in programming of the offspring metabolic, immune and nervous system development has been well documented in humans and food-producing animals including cattle (Palmer, 2011; Caton et al., 2019), the potential involvement of the maternal microbiome in the developmental origins of health and disease has recently began to be better appreciated (Stiemsma and Michels, 2018; Calatayud et al., 2019; Codagnone et al., 2019). It was hypothesized that undesired alterations in the maternal microbiota could indirectly influence fetal development, and that these effects may subsequently be transmitted to progeny, resulting in the development of an altered microbiota in offspring (Calatayud et al., 2019). Undesirable outcomes in offspring resulting from changes in the maternal microbiota include increased susceptibility to the development of metabolic disorders and respiratory infections (Calatayud et al., 2019; Yao et al., 2020). Recent evidence from studies in mice demonstrated that the maternal microbiota during pregnancy modulates the programming of fetal metabolic and nervous system development (Kimura et al., 2020; Vuong et al., 2020). Considering the increased evidence showing the importance of the maternal microbiota in developmental programming in rodent models, and the greater evidence regarding the involvement of the microbiome in defining cattle health and productivity, exploring the role of the maternal microbiota in fetal programming and offspring microbiome development may provide important information for improving cattle health and feed efficiency.

In the present study, we used 16S rRNA gene sequencing to characterize the nasopharyngeal, ruminal and vaginal microbiota of virgin yearling heifers from dams given different nutritional diets during their first trimester of gestation, and in pregnant beef heifers in response to direct feeding of a mineral and vitamin (VTM) supplement during the first 6 months of gestation. Of note, a well-defined positive impact of maternal VTM supplementation exists on offspring health and performance in beef cattle, and the role of VTM on fetal programming assessed during the first trimester of pregnancy have been documented (Mee et al., 1995; Wilde, 2006; Van Emon et al., 2020; Diniz et al., 2021; Menezes et al., 2021). Questions remain, however, pertaining to whether these maternal VTM supplementation-associated positive outcomes are dependent on VTM-induced alterations of ruminal microbiota. To provide a more holistic view of the microbiota residing within the respiratory, gastrointestinal, and reproductive tract of cattle, the similarity of the microbiota within these sites was evaluated, and taxa shared amongst the three microbial habitats were identified. Given the relevance of these microbial communities to respiratory and reproductive health and rumen fermentation/nutrient metabolism, and most importantly, as potential maternal inoculant sources for seeding the fetal and offspring microbiota, a holistic evaluation of bovine microbiota is therefore necessary rather than focusing on only one microbial community.

## MATERIALS AND METHODS

Animals used in this study were cared for in accordance with the guidelines set by the Olfert et al. (1993) and all experimental procedures involving cattle were approved by the North Dakota State University Institutional Animal Care and Use Committee (#A20085 and #A20085, for virgin yearling heifers and for pregnant heifers, respectively).

### Animal Husbandry and Experimental Design

#### Virgin yearling heifers

Deep nasopharyngeal swabs, ruminal fluid, and vaginal swabs were collected from 45 F1 virgin heifers (9-month-old, BW = 688 ± 57 kg) whose dams were assigned to either a low gain treatment (**LG**, targeted average daily gain of 0.28 kg/d, n = 22) or a moderate gain treatment (**MG**, 0.79 kg/d, n = 23) during the first 84 days of gestation. To achieve the LG, dams were maintained on a basal diet consisting of prairie grass hay, corn silage, and dried distillers grains plus solubles. To achieve the MG (0.79 kg/d), heifers were fed the basal diet with the addition of a protein/energy supplement fed at the rate of 0.58% BW as-fed daily. Up to d 84 of gestation dams were housed and individually fed (Insentec; Hokofarm B.V. Repelweg 10, 8316 PV Marknesse, the Netherlands) at the Beef Cattle Research and Extension Center (BCRC) in Fargo, ND. After day 84 of gestation, dams were transported to the Central Grasslands Research Extension Center (CGREC) in Streeter, ND, where they were managed as a single group on common diets until parturition and subsequent weaning of the F1 offspring. Upon weaning, the F1 heifers (approx. 6-months old) were transported to the BCRC where they were housed in two pens and individually fed (Insentec; Hokofarm B.V. Repelweg 10, 8316 PV Marknesse, the Netherlands) a common diet (Table 1).

**Table 1.** Nutrient composition of the diets fed to the dams of virgin yearling heifers during the first 84 days of gestation, and virgin yearling heifers and pregnant heifers at the time of sample collection.

| Diet composition, % DM | Dams of virgin yearling |  | Virgin yearling | Pregnant heifers |  |
| --- | --- | --- | --- | --- | --- |
|  | LG<sup>1</sup> | MG<sup>2</sup> |  | Control | VTM<sup>3</sup> |
| Corn silage | 37 | 29 | 20 | 30 | 30 |
| Prairie hay | 53 | 41 | 70 | 59 | 59 |
| Dried distillers grains plus solubles | 10 | 5 | 5 | 6 | 6 |
| Premix | - | - | 5 | 5 | 5 |
| Energy and protein supplement<sup>1</sup> | - | 25 | - | - | - |

<sup>1</sup>Basal total mixed rations (TMR) contained a commercially available mineral supplement (Purina® Wind & Rain® Storm® All-Season 7.5 Complete Mineral, Land O'Lakes Inc., Arden Hills, Minn.) fed at a rate of 113.4 gram per head per day, targeting gain of 0.28 kg/d.
<sup>2</sup>The supplement fed was an energy/protein supplement formulated with a blend of ground corn, DDGS, wheat midds, fish oil and urea, targeting gain of 0.79 kg/d.
<sup>3</sup>VTM: Vitamin mineral supplement was a pelleted product fed at 0.45 kg/head/day (consisting of 113 g of a vitamin and mineral supplement [Purina Wind & Rain Storm All-Season 7.5 Complete, Land O'Lakes Inc., Arden Hills, Minn.] and 337 g of a carrier) (Menezes et al., 2021).

#### Pregnant heifeirs

The nasopharyngeal, ruminal, and vaginal microbiota of the replacement pregnant heifers (1 year 9 months old, BW = 1001 ± 128 kg) during the sixth month of gestation were also evaluated. At breeding, heifers were assigned to one of two treatments: 1) vitamin and mineral supplementation (**VTM**; n = 17) or 2) no vitamin and mineral supplementation (**Control**; n = 15). Heifers were housed at the BCRC and individually fed (Insentec; Hokofarm B.V. Repelweg 10, 8316 PV Marknesse, the Netherlands) a total mixed ration containing triticale hay, corn silage, modified distillers grains plus solubles, ground corn, and, if indicated by treatment, a VTM premix (Table 1). The VTM premix was fed at 0.45 kg/heifer/day to provide macro and trace minerals and vitamins A, D, and E to meet 110% of the daily requirements (NASEM, 2016). The specific ingredients within the VTM supplement are as previously described (Menezes et al., 2021).

### Nasopharyngeal swab, ruminal fluid, and vaginal swab sampling

Nasopharyngeal swabs, ruminal fluid and vaginal swabs were collected simultaneously from each of the virgin yearling and pregnant heifers by same personnel on the same day. All sample collection was completed within 4 hours.

#### Nasopharyngeal sampling

Deep nasopharyngeal swabs were collected as previously described (Holman et al., 2017; Amat et al., 2019). Briefly, prior to sampling, the right nostril of the heifer was wiped clean with 70% ethanol and an extended guarded swab (27 cm) with a rayon bud (MW 128, Medical Wire & Equipment, Corsham, England) was used for sampling. Swab tips were then be cut and placed in a sterile 2 mL centrifuge tube on ice until processing. Upon arrival in the lab, the swab was transferred into 1 mL sterile brain heart infusion (BHI) broth containing 20% glycerol stock.

#### Rumen fluid sampling

Rumen fluid sample collection was performed using the method currently used in our laboratory which was modified from Paz et al. (2016). Briefly, a rigid metal speculum was placed into the mouth of the heifer and then a flexible plastic tube with multiple holes at the tip was passed through the speculum and into the esophagus. The speculum was used to ensure that the plastic tube was not damaged by the heifers and that the tube entered the esophagus. Once the tube entered the rumen, and was below the ruminal mat layer, suction pressure was applied to the tube to collect ruminal fluid. Up to 120 mL of ruminal fluid was collected on each sampling day. Separate tubing and containers were used for each heifer to avoid cross-contamination. After thorough mixing, an aliquot of 40 ml of rumen fluid was placed into a 50 mL falcon tube and immediately frozen with dry ice.

#### Vaginal sampling

For vaginal sampling, the vulva was thoroughly cleaned with 70% ethanol and a paper towel. Then the labia majora of the vulva was held open allowing the passage of a swab (15 cm, sterile cotton tipped applicators with aerated tip protector; Puritan). When the swab tip reached the midpoint of the vaginal cavity, it was placed against the vaginal wall, swirled four times, and then withdrawn carefully to minimize contamination. The vaginal swabs were immediately placed in sterile Whirl Pak bags and transported on ice to the lab. All nasopharyngeal and vaginal swabs as well as rumen fluid were stored at -80°C until DNA extraction.

### Metagenomic DNA extraction

Metagenomic DNA was extracted from the nasopharyngeal and vaginal swabs using a Qiagen DNeasy Tissue kit (Qiagen Inc., Germantown, MD, USA) according to the kit manual with minor modifications. Briefly, the cotton tip of the nasopharyngeal swab was removed and placed back into the BHI-glycerol mixture, and then centrifuged at 20,000 × g for 5 min at 4°C to pellet the cotton and microbes. The pellet was then re-suspended in 180 μl of enzymatic buffer. The enzymatic buffer [20 mM Tris.CI (pH 8.0), 2mM sodium EDTA, and 1.2% Triton X-100] contained 300 U/ml mutanolysin and 20 mg/ml lysozyme. The mixture was then vortexed and incubated for 1 h at 37°C with agitation at 800 rpm. After incubation, 25 μl proteinase K and 200 μl Buffer AL (without ethanol) were added and vortexed, and then incubated at 56°C for 30 min with agitation at 800 rpm. Approximately 400 mg of 0.1 mm zircon/silica beads were added to the tube and subjected to mechanical cell lysis using a FastPrep-24 Classic bead beater (MP Biomedicals, Irvine, CA) at 4.0 m/s for 24 s. The mixture was then centrifuged (13,000 × g for 5min), and the supernatant (approx. 300-400 µl) was removed and placed in a new tube and mixed with an equal volume of 100% ethanol. From this step onward, the procedures were performed as described in the DNeasy Tissue Kit instruction manual.

The procedures for metagenomic DNA extraction from the vaginal swabs were identical to those used on the nasopharyngeal swabs with the following exceptions. First, the cotton swab was removed from applicator and placed in a sterile 2 mL centrifuge tube. Then, 360 µl of enzymatic buffer was added to the tube to ensure complete emersion of the swab in the enzymatic buffer. Metagenomic DNA from the rumen fluid samples was extracted using the Qiagen DNeasy PowerLyzer PowerSoil kit (Qiagen Inc.) according to the instructions of manufacturer. The frozen rumen fluid samples were thawed, and vortexed thoroughly before transfer to a sterile 2 mL microfuge tube. The sample was then centrifuged at 20,000 × *g* for 10 min at 4°C to pellet the microbes in the sample. From this point onwards, the protocol for the PowerLyzer PowerSoil kit was followed as per the instructions of the manufacturer. Negative extraction controls were included for all extraction kits.

### 16S rRNA gene sequencing and analysis

The V3-V4 hypervariable regions of the 16S rRNA gene were amplified using the 341-F (5’-CCTAYGGGRBGCASCAG-3’) and 806-R (5’-GGACTACNNGGGTATCTAAT-3’) primers. All PCR steps were carried out using the Phusion High-Fidelity PCR Master Mix (New England Biolabs). The PCR products were electrophoresed on a 2% agarose gel and stained with SYBR Safe DNA gel stain. The DNA fragment was excised from the gel and purified using the QIAquick Gel Extraction Kit (Qiagen Inc.,). Sequencing libraries were generated with NEBNext Ultra DNA Library Prep Kit (New England BioLabs, Ipswich, MA, USA) for Illumina, following the recommendations of the manufacturer. The library quality was assessed with a Qubit 2.0 Fluorometer (Thermo Scientific) and Agilent Bioanalyzer 2100 system. Libraries were then sequenced on a NovaSeq 6000 instrument with a SP flow cell (2 x 250 bp) (Illumina Inc., San Diego, CA, USA).

The 16S rRNA gene sequences were processed using DADA2 v. 1.18 (Callahan et al., 2016) in R. 4.0.3. Briefly, the forward reads were truncated at 225 bp and the reverse reads at 220 bp. The reads were merged, chimeric sequences removed, and taxonomy assigned to each merged sequence, referred to here as operational taxonomic units (OTUs) at 100 % similarity, using the naïve Bayesian RDP classifier (Wang et al., 2007) and the SILVA SSU database release 138 (Quast et al., 2013). OTUs that were predominantly in the negative extraction control samples and likely to be contaminants were removed prior to analyses as were those OTUs classified as chloroplasts, mitochondria, or eukaryota. The number of OTUs per sample (richness), the Shannon and inverse Simpson’s diversity indices, and Bray-Curtis dissimilarities were calculated in R using Phyloseq 1.34.0 (McMurdie and Holmes, 2013) and vegan 2.5-7 (Oksanen et al., 2013). To account for uneven sequence depths, samples were randomly subsampled to 7,100, 73,500, and 10,300 for the nasopharyngeal, ruminal, and vaginal samples respectively, prior to the calculation of Bray-Curtis dissimilarities and diversity measures for the virgin heifers. For the pregnant heifers, these values were 6,200, 74,500, and 8,200.

### Statistical Analysis

Permutational multivariate analysis of variance (PERMANOVA; adonis2 function; 10,000 permutations) of the Bray-Curtis dissimilarities was performed using vegan to determine the effect of maternal nutrition on the nasopharyngeal, ruminal and vaginal microbial community structure in virgin heifers whose dams were managed to targeted LG or MG during the first 84 days of gestation. The effect of VTM supplementation on the microbial community structure of these three microbiotas in pregnant heifers was also assessed. Differentially abundant genera between treatment groups for both the virgin and pregnant heifers were identified using MaAsLin2 v. 1.5.1 in R (Mallick et al., 2021). Only those genera with a relative abundance greater than 0.1% within each sample type were included. Diversity metrics were compared by treatment for both virgin and pregnant heifers using an unpaired *t*-test. The number of OTUs (richness), diversity indices, relative abundance of the most relatively abundant genera between the LG and MG groups of virgin yearling heifers, or between the VTM and Control groups of pregnant heifers, and the relative abundance of *Methanobrevibacter* spp. between the virgin and pregnant heifers were compared using the generalized liner mixed model estimation procedure (PROC GLIMMIX) in SAS (ver. 9.4, SAS Institute Inc. Cary, NC). The means were compared using the LSMEANS statement and significance was declared at *P* < 0·05.

Spearman’s rank-based correlations between *Methanobrevibacter* and the other 24 most relatively abundant genera in the ruminal and vaginal microbiota were calculated using the CORR procedure in SAS with the SPEARMAN option. From these 24 genera, the genera that have significant effect on *Methanobrevibacter* abundance were predicted using a stepwise-selected GLIMMIX model with beta-binomial distribution as described previously (Amat et al., 2019). The model used was: logit (Ŷ) = ln (π/(1 – π)) = b0 + b1 (X1) + … + bn (Xn), where π represents the relative abundance of the *Methanobrevibacter* genus (0–1) and Xn represents the relative abundance (0–100%) of a bacterial genus n. The stepwise selection method involved backward elimination and forward selection to eliminate any variables in the model that have no significant effect (*p* > 0.05) on the predicted outcome.

## RESULTS

### Sequencing Results

An average of 66,045 ± 31,588 (SD) 16S rRNA gene sequences per sample (min. = 2,374; max. = 139,012) were obtained from 219 nasopharyngeal, ruminal fluid, and vaginal samples. From these sequences, a total of 81,391 archaeal and bacterial OTUs were identified and classified into 58 unique phyla (8 archaeal and 50 bacterial phyla), and 1,511 unique genera.

### Effect of Maternal Restricted Gain During the First Trimester of Gestation on Offspring Microbiota Development

To determine the effect of maternal nutrition during the first trimester of gestation on microbial populations of their offspring, we characterized and compared the nasopharyngeal, ruminal and vaginal microbiota of virgin yearling heifers from two groups of dams that were subjected to diets resulting in either a LG or MG phenotype during the first 84 days of gestation. The microbial community structure of the nasopharynx (PERMANOVA: R^2^ = 0.027, *p* = 0.57), rumen (PERMANOVA: R^2^ = 0.02, *p* = 0.98) and vagina (PERMANOVA: R^2^ = 0.028, *p* = 0.37) in the virgin heifers did not differ between the LG and MG groups (Fig. 1A). Microbial richness and diversity as measured by the number of OTUs, and the Shannon and inverse Simpson’s diversity indices of these microbiotas also did not significantly differ by maternal nutrition group (Fig. 1B, C and D; *p* > 0.05). There was, however, a strong tendency (*p* = 0.06) in LG offspring to harbor a richer ruminal microbial community compared to MG offspring (2605 vs. 2515 OTUs).

**Figure 1.**
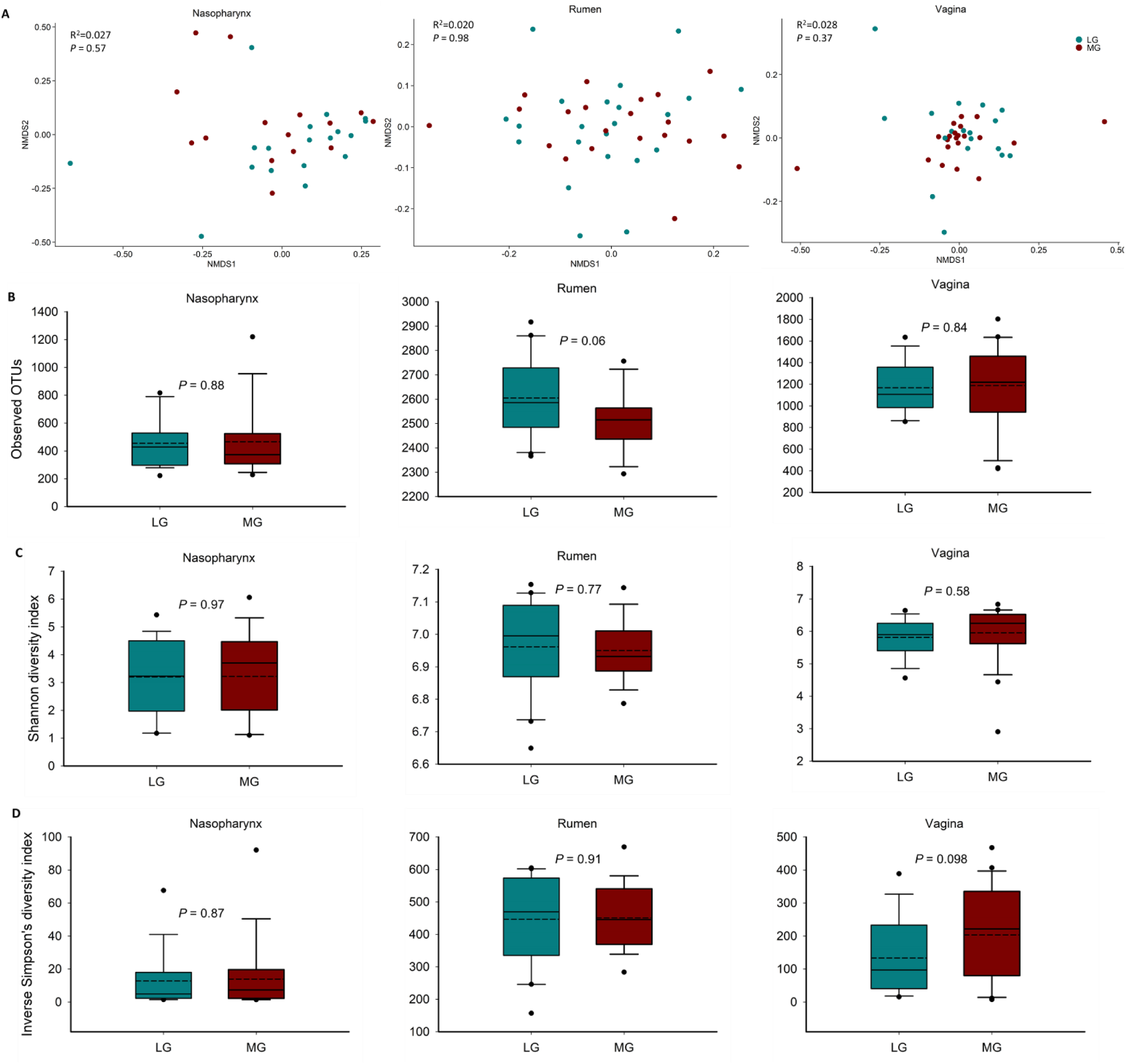
Beta and alpha diversity of the nasopharyngeal, ruminal and vaginal microbiota of virgin yearling heifers from low gain (LG) or medium gain (MG) dams as determined during the first trimester of gestation. (**A**) Non-metric multidimensional scaling (NMDS) plots of the Bray-Curtis dissimilarities, (**B**) number of operational taxanomic units (OTUs), and Shannon (**C**) and inverse Simpson’s diversity index (**D**) of each microbial community.

The nasopharyngeal microbiota across all animals was dominated by *Actinobacteriota* (51%), *Firmicutes* (28.2%), *Bacteroidota* (10.8%) and *Proteobacteria* (4.9%). The relative abundance of the eight most relatively abundant phyla did not differ between LG and MG virgin heifers (*p* > 0.05) (Fig. 2A). *Bacteroidota* was the most relatively abundant phylum in the ruminal microbiota (65.5%) followed by *Firmicutes* (24.2%). As with the nasopharyngeal microbiota, none of the eight most relatively abundant phyla in the rumen microbiota differed between the two treatment groups. The most relatively abundant phylum present in the vaginal tract was *Firmicutes* (52%) followed by *Bacteroidota* (23.0%) and *Actinobacteriota* (17.4%). Similar to the rumen and nasopharyngeal microbiota, no difference between treatments was detected in the relative abundance of eight most relatively abundant phyla in the vaginal microbiota (*p* > 0.05).

**Figure 2.**
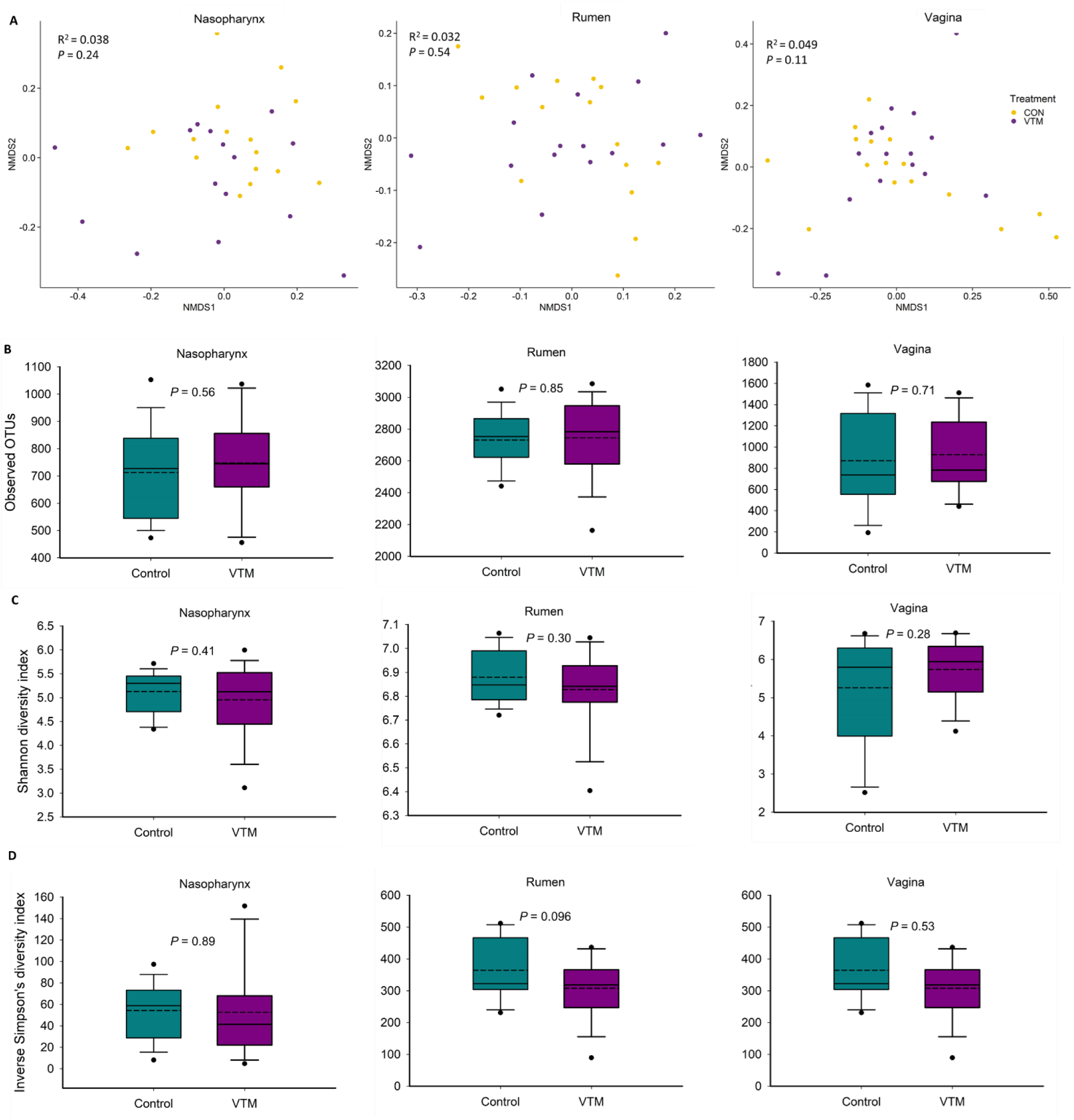
Beta and alpha diversity of the nasopharyngeal, ruminal and vaginal microbiota of pregnant heifers that received a vitamin and mineral supplement (VTM) or a control diet (Control) during the first six months of gestation. **A**) Non-metric multidimensional scaling (NMDS) plots of the Bray-Curtis dissimilarities, **B**) number of operational taxanomic units (OTUs), and Shannon (**C**) and inverse Simpson’s diversity index (**D**) of each microbial community.

The 25 most relatively abundant genera in the nasopharyngeal, ruminal and vaginal microbiota are listed in Table 2. Overall, the predominant nasopharyngeal genera did not differ in their relative abundance between the LG and MG groups (*p* > 0.05). In the rumen, the relative abundance of only one genus ([*Eubacterium] ruminantium group)* was significantly different between the two groups, being greater in the MG group than in the LG group (*p* = 0.029). Within the vaginal microbial community, *Alistipes*, *Ruminococcus* and *Oscillospiraceae* NK4A214 group were significantly less relatively abundant in the LG group compared to the MG group (*p* < 0.05). The relative abundance of *Romboutsia* (*p* = 0.061) and *Paeniclostridium* (*p* = 0.092) tended to be lower while *Arcanobacterium* (*p* = 0.090) tended to be higher in MG group compared to LG group.

**Table 2.** Percent relative abundance of the most relatively abundant genera in nasopharyngeal (n = 26), ruminal (n= 24) and vaginal (n = 27) microbiota of virgin yearling heifers that were born from the dams received a basal diet to achieve a moderate gain (MG) or to achieve a low gain (LG) during the first 84 days of gestation^1^

| Nasopharynx |  |  |  |  |  | Rumen |  |  |  |  |  | Vagina |  |  |  |  |  |
| --- | --- | --- | --- | --- | --- | --- | --- | --- | --- | --- | --- | --- | --- | --- | --- | --- | --- |
| Genus | Rank | MG | LG | SEM | P-value | Genus | Rank | MG | LG | SEM | P-value | Genus | Rank | MG | LG | SEM | P-value |
| *Mycoplasma* | 2 | 23.8 | 12.3 | 9.16 | 0.218 | *Prevotella* | 1 | 31.4 | 30.2 | 1.85 | 0.505 | *Corynebacterium* | 1 | 8.75 | 12.00 | 2.54 | 0.209 |
| *Cellulomonas* | 3 | 5.41 | 6.40 | 1.92 | 0.608 | *Rikenellaceae RC9 gut group* | 2 | 8.89 | 9.14 | 0.84 | 0.769 | *Oscillospiraceae UCG-005* | 2 | 9.52 | 7.86 | 0.91 | 0.076 |
| *Filobacterium* | 4 | 2.62 | 8.03 | 3.00 | 0.081 | *Christensenellaceae R-7 group* | 4 | 4.70 | 4.81 | 0.48 | 0.804 | *Rikenellaceae RC9 gut group* | 4 | 4.30 | 3.87 | 0.51 | 0.406 |
| *Arthrobacter* | 5 | 2.36 | 2.05 | 0.51 | 0.546 | *Methanobrevibacter* | 5 | 4.30 | 4.47 | 0.55 | 0.756 | *Bacteroides* | 5 | 4.15 | 3.95 | 0.62 | 0.757 |
| *Corynebacterium* | 6 | 1.60 | 1.81 | 0.45 | 0.636 | *Prevotellaceae UCG-003* | 7 | 3.82 | 3.77 | 0.19 | 0.803 | *Christensenellaceae R-7 group* | 8 | 3.25 | 3.22 | 0.37 | 0.952 |
| *Nocardioides* | 7 | 1.77 | 1.65 | 0.60 | 0.842 | *Prevotellaceae UCG-001* | 8 | 2.77 | 2.85 | 0.19 | 0.660 | *Alistipes* | 9 | 3.55 | 2.55 | 0.40 | 0.017 |
| *Oscillospiraceae UCG-005* | 9 | 1.35 | 1.36 | 0.58 | 0.975 | *Ruminococcus* | 9 | 2.55 | 2.36 | 0.18 | 0.305 | *Monoglobus* | 10 | 2.83 | 2.44 | 0.29 | 0.182 |
| *Streptomyces* | 13 | 1.06 | 0.76 | 0.34 | 0.389 | *Fibrobacter* | 12 | 1.36 | 1.45 | 0.30 | 0.773 | *Romboutsia* | 11 | 2.44 | 1.89 | 0.28 | 0.061 |
| *Romboutsia* | 15 | 0.85 | 0.80 | 0.22 | 0.809 | *Oscillospiraceae NK4A214 group* | 14 | 1.42 | 1.34 | 0.10 | 0.444 | *Prevotellaceae UCG-003* | 13 | 1.71 | 1.79 | 0.29 | 0.771 |
| *Christensenellaceae R-7 group* | 16 | 0.83 | 0.71 | 0.31 | 0.702 | *Papillibacter* | 18 | 1.11 | 1.17 | 0.13 | 0.668 | *Campylobacter* | 16 | 1.24 | 1.69 | 1.03 | 0.668 |
| *Ornithinimicrobium* | 18 | 0.55 | 0.72 | 0.31 | 0.600 | *Anaeroplasma* | 21 | 0.96 | 0.90 | 0.10 | 0.551 | *Trueperella* | 17 | 0.66 | 2.30 | 1.14 | 0.160 |
| *Rikenellaceae RC9 gut group* | 19 | 0.49 | 0.56 | 0.18 | 0.697 | *Pseudobutyrvibrio* | 22 | 0.82 | 0.81 | 0.05 | 0.856 | *Ruminococcus* | 18 | 1.64 | 1.20 | 0.19 | 0.025 |
| *Bacteroides* | 21 | 0.52 | 0.49 | 0.16 | 0.865 | *Prevotellaceae NK3B31 group* | 26 | 0.69 | 0.79 | 0.08 | 0.193 | *Lachnospiraceae NK3A20 group* | 21 | 1.07 | 0.90 | 0.17 | 0.307 |
| *Marmoricola* | 22 | 0.58 | 0.41 | 0.13 | 0.187 | *Lachnospiraceae NK3A20 group* | 30 | 0.47 | 0.45 | 0.05 | 0.599 | *Paeniclostridium* | 22 | 1.13 | 0.79 | 0.19 | 0.092 |
| *Ruminococcus* | 24 | 0.43 | 0.50 | 0.18 | 0.658 | *Lachnospiraceae XPB1014 group* | 31 | 0.44 | 0.44 | 0.05 | 0.957 | *Arcanobacterium* | 24 | 0.49 | 1.35 | 0.50 | 0.090 |
| *Lachnospiraceae NK3A20 group* | 25 | 0.53 | 0.31 | 0.14 | 0.110 | *Prevotellaceae UCG-004* | 32 | 0.44 | 0.41 | 0.05 | 0.547 | *Family XIII AD3011 group* | 25 | 0.97 | 0.81 | 0.10 | 0.116 |
| *Prevotella* | 26 | 0.58 | 0.24 | 0.26 | 0.196 | *Lachnospiraceae ND3007 group* | 33 | 0.39 | 0.44 | 0.06 | 0.429 | *Streptobacillus* | 26 | 1.58 | 0.07 | 1.00 | 0.139 |
| *Ornithinococcus* | 29 | 0.33 | 0.42 | 0.13 | 0.527 | *Rikenellaceae U29-B03* | 34 | 0.40 | 0.41 | 0.07 | 0.997 | *Akkermansia* | 27 | 0.85 | 0.79 | 0.20 | 0.754 |
| *Clostridium sensu stricto 1* | 30 | 0.41 | 0.34 | 0.14 | 0.615 | *[Eubacterium] hallii group* | 36 | 0.40 | 0.38 | 0.03 | 0.573 | *Porphyromonas* | 28 | 0.34 | 1.23 | 1.01 | 0.382 |
| *Olsenella* | 31 | 0.32 | 0.39 | 0.14 | 0.618 | *Butyrivibrio* | 38 | 0.38 | 0.33 | 0.05 | 0.295 | *Butyricoccaceae UCG-009* | 30 | 0.74 | 0.66 | 0.09 | 0.395 |
| *Bifidobacterium* | 32 | 0.33 | 0.37 | 0.17 | 0.819 | *Succinivibrio* | 39 | 0.32 | 0.36 | 0.04 | 0.402 | *Oscillospiraceae NK4A214 group* | 31 | 0.81 | 0.56 | 0.09 | 0.010 |
| *Paeniclostridium* | 33 | 0.40 | 0.31 | 0.11 | 0.393 | *Prevotellaceae YAB2003 group* | 40 | 0.35 | 0.32 | 0.04 | 0.435 | *Rikenellaceae dgA-11 gut group* | 33 | 0.71 | 0.59 | 0.08 | 0.149 |
| *Prevotellaceae UCG-003* | 35 | 0.43 | 0.22 | 0.13 | 0.111 | *[Eubacterium] ruminantium group* | 41 | 0.32 | 0.27 | 0.02 | 0.029 | *Prevotellaceae UCG-004* | 37 | 0.68 | 0.54 | 0.11 | 0.209 |
| *Monoglobus* | 36 | 0.30 | 0.32 | 0.13 | 0.859 | *Olsenella* | 42 | 0.27 | 0.26 | 0.04 | 0.754 | *Lachnospiraceae UCG-010* | 38 | 0.62 | 0.59 | 0.08 | 0.683 |
| *Rhodococcus* | 37 | 0.28 | 0.31 | 0.12 | 0.778 |  |  |  |  |  |  | *Clostridium sensu stricto 1* | 39 | 0.54 | 0.68 | 0.21 | 0.514 |
| *Brachybacterium* | 38 | 0.31 | 0.27 | 0.12 | 0.777 |  |  |  |  |  |  | *Olsenella* | 40 | 0.30 | 0.92 | 0.55 | 0.263 |
| *Mannheimia\** |  | 0.01 | 0.00 |  |  | *Fusobacterium\** |  | 0.01 | 0.02 |  |  | *Alloprevotella* | 42 | 0.57 | 0.42 | 0.13 | 0.246 |
| *Pasteurella\** |  | 0.02 | 0.07 |  |  | *Trueperella\** |  | 0.00 | 0.00 |  |  | *Fusobacterium\** |  | 0.00 | 0.00 |  |  |
| *Histophilus\** |  | 0.00 | 0.01 |  |  |  |  |  |  |  |  |  |  |  |  |  |  |

<sup>1</sup>The genera whose relative abundance was ranked within the top 42 are listed in this table and any ones within top 42 rank that were unclassified at genus level were excluded.
\*These genera included in this table because of their relevance to bovine respiratory disease and liver abscesses in cattle.

### Effect of Vitamin and Mineral Supplementation During the First Six Months of Gestation on the Maternal Microbiota

To investigate whether VTM supplementation during the first 6 months of gestation affects the maternal microbiota, we compared the nasopharyngeal, ruminal and vaginal microbiota between the VTM and Control groups of replacement pregnant heifers. The community structure of the nasopharyngeal (PERMANOVA: R^2^ = 0.038, *p* > 0.05), ruminal (PERMANOVA: R^2^ = 0.032, *p* > 0.05) and vaginal (PERMANOVA: R^2^ = 0.049, *p* > 0.05) microbiota was not affected by VTM supplementation (Fig.2A). Microbial richness and diversity also did not differ by VTM supplementation in any of the three microbial communities (*p* > 0.05) (Fig. 2B, C and D). At the phylum level, the relative abundance of the eight most relatively abundant phyla in the nasopharynx, rumen and vagina did not differ between the control and VTM groups (Fig. 3B). However, the relative abundance of several genera present in the nasopharynx (5 genera) and rumen (3 genera) was affected by VTM supplementation (*p* < 0.05, Table 3).

**Figure 3.**
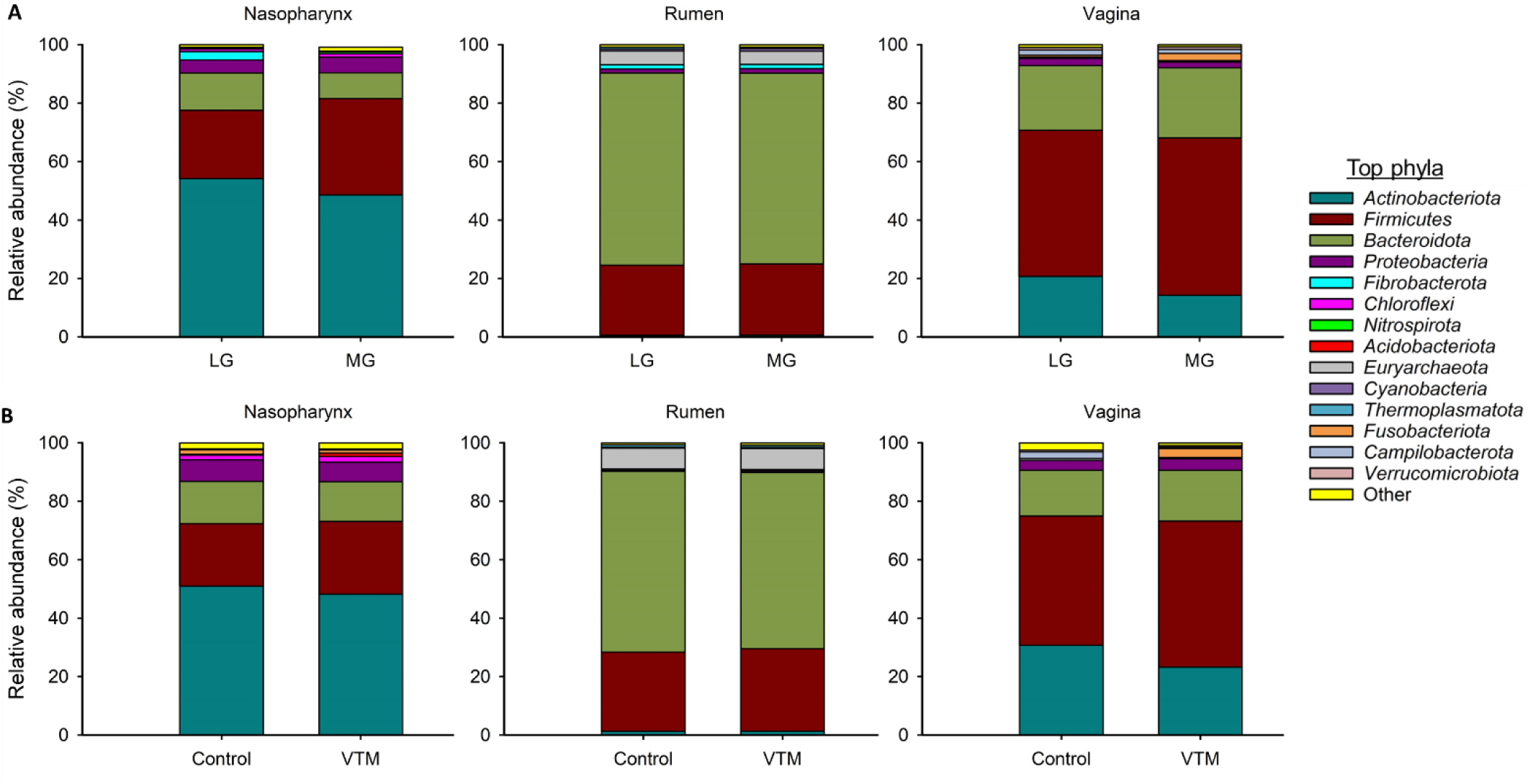
Percent relative abundance of the eight most relatively abundant phyla in the nasopharyngeal, ruminal and vaginal microbiota of (**A**) virgin yearling heifers from low gain (LG) or medium gain (MG) dams and (**B**) pregnant heifers that received a vitamin and mineral supplement (VTM) or a control diet (Control) during the first 6 six months of gestation

**Table 3.**
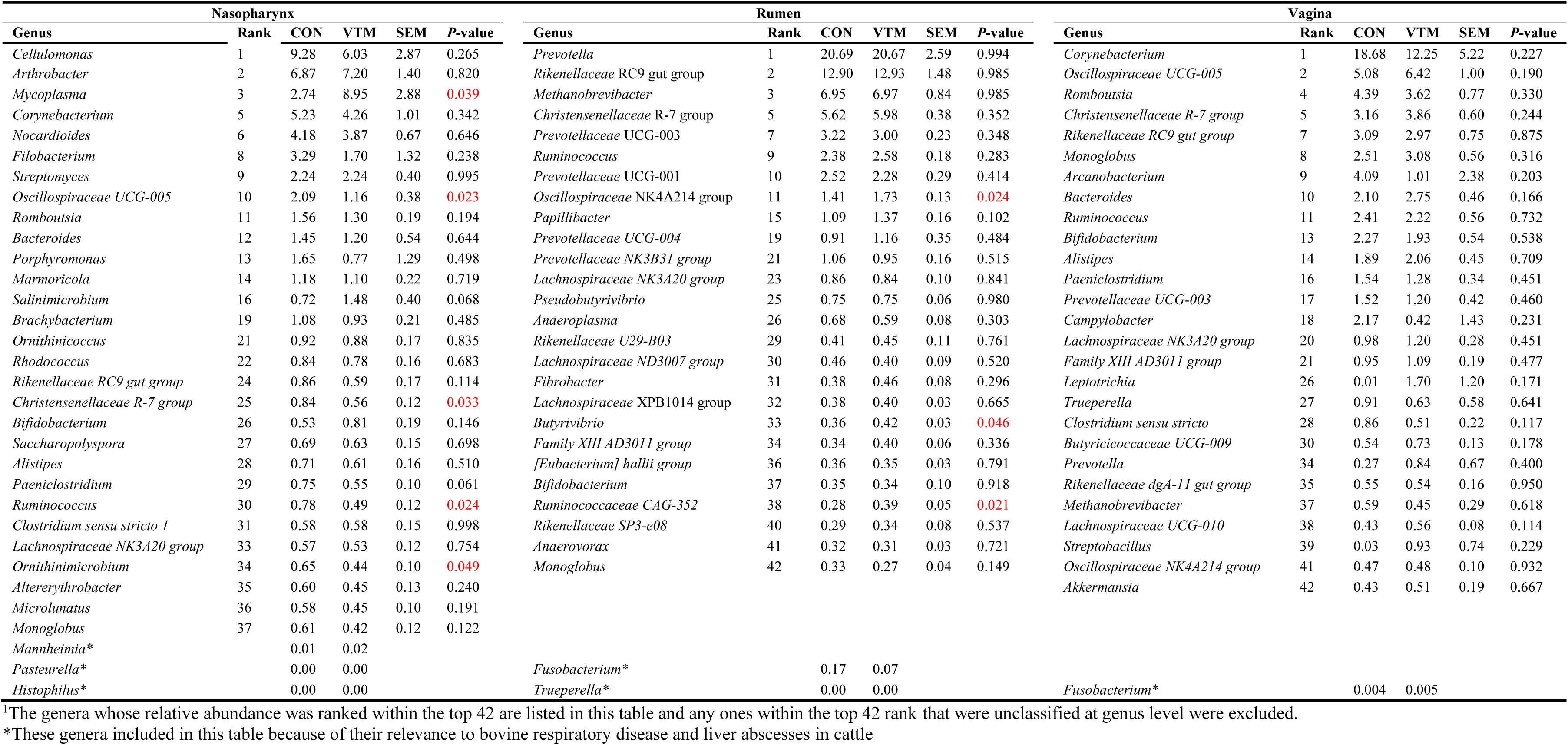
Percent relative abundance of the 42 most relatively abundant genera in nasopharyngeal (n = 29), ruminal (n= 26) and vaginal (n = 27) microbiota of received diets without (CON) and with vitamin and mineral supplementation (VTM) during the first 6 months of gestation^1^

*Mycoplasma*, the third most relative abundant genera in the nasopharyngeal microbiota, was enriched in pregnant heifers receiving the VTM supplement (8.95% vs. 2.74%, *p* = 0.039). VTM supplementation also resulted in a reduced relative abundance of *Oscillospiraceae* UCG-005, *Christensenellaceae R-7 group*, *Ruminococcus* and *Ornithinimicrobium* genera (*p* < 0.05). Among the 26 most predominant ruminal genera, statistically significant difference in relative abundance was observed in only three genera (*Oscillospiraceae* NK4A214 group, *Butyrivibrio* and *Ruminococcaceae* CAG-352), and all of which were enriched in the VTM group (*p* < 0.05). The relative abundance of the 27 relatively most abundant genera in the vaginal microbiota did not differ between the VTM and control groups (*p* > 0.05) (Table 3).

### Holistic View of Nasopharyngeal, Ruminal and Vaginal Microbiota and Identification of Core Taxa Shared Across These Microbiomes

To provide a holistic view of the microbiota residing within the respiratory, gastrointestinal, and reproductive tracts of each animal, we attempted to identify similarities among these microbial communities in virgin and pregnant heifers. To do this, the sequence data from all animal groups and treatments were combined and all samples were randomly subsampled to 6,200 sequences. As expected, each anatomical site had a distinct microbiota (Fig.4). In terms of microbial richness, the rumen had the richest microbiota followed by the vagina and nasopharynx in both virgin (Fig.1B) and pregnant heifers (Fig. 2B). Overall, the ruminal and vaginal microbiota were also more diverse than the nasopharyngeal microbiota in both groups of heifers (Fig. 1C and D and Fig. 2C and D).

**Figure 4.**
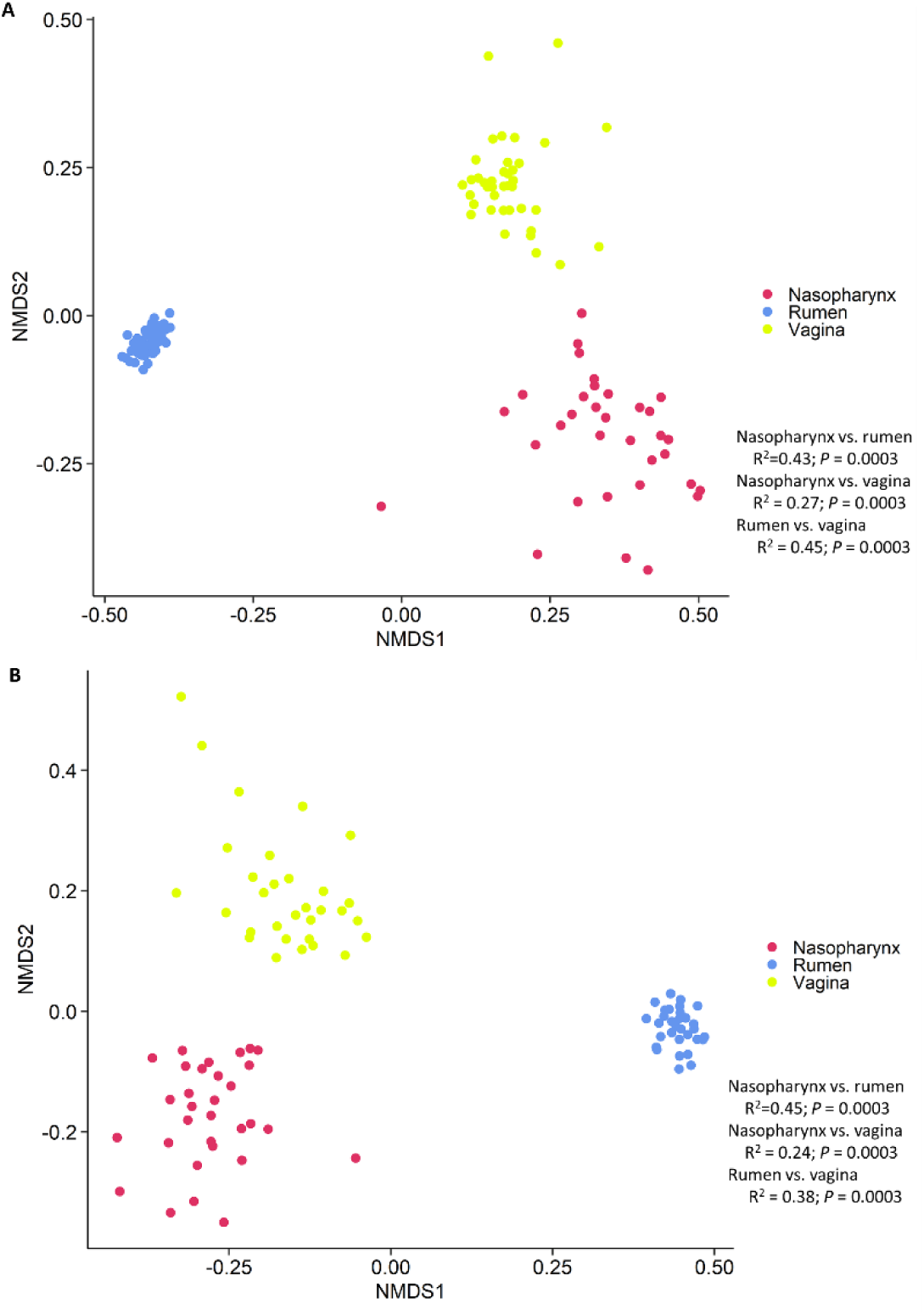
Non-metric multidimensional scaling (NMDS) plots of the Bray-Curtis dissimilarities of the nasopharyngeal, ruminal and vaginal microbiota of (**A**) virgin yearling and (**B**) pregnant heifers.

Many taxa appeared to be highly specific to one of the three microbial habitats as shown in the heatmap (Fig. 5). For example, OTUs classified as *Prevotella*, *Papillibacter*, *Oscillospiraceae NK4A214* group and *Pseudobutyrivibrio* were more exclusively present in the rumen. While the most of the OTUs within the archaeal *Methanobrevibacter* genus were present in all three habitats, the rumen was most predominantly colonized by members of this genera. Some taxa, including *Mycoplasma*, *Filobacterium*, *Streptomyces*, *Nocardioides*, *Marmoricola*, *Arthrobacter* and *Cellulomonas spp.*, were associated with the nasopharynx. Certain *Corynebacterium* OTUs appeared to be specific to the vaginal microbiota.

**Figure 5.**
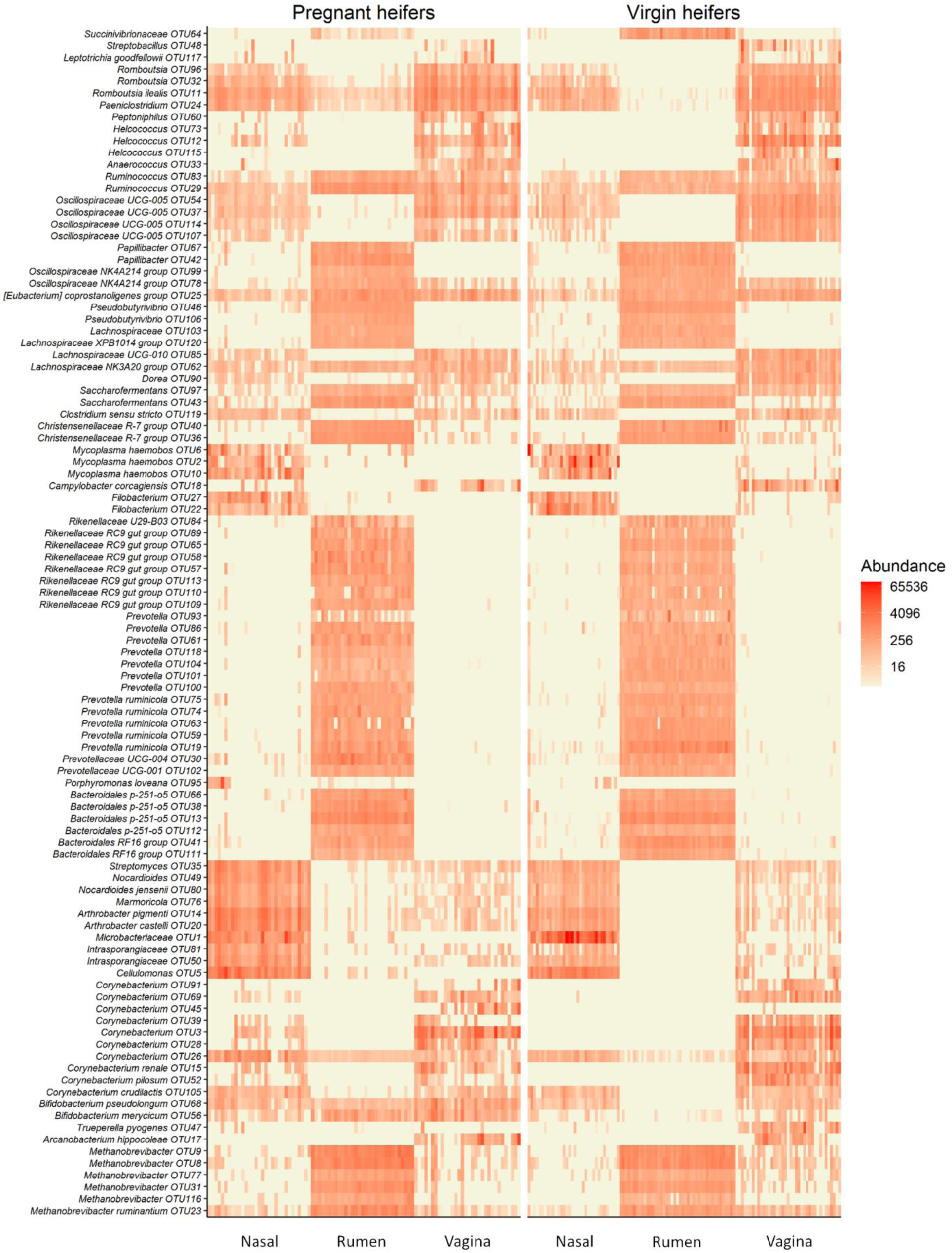
Heat map showing the 100 most abundant OTUs (log_4_) overall by sample type within each animal group (Pregnant and virgin heifers).

Although the nasopharyngeal, ruminal and vaginal microbiota were clearly distinct, a small number of taxa were present in all three microbial communities. We identified 43 OTUs that were shared by more than 60% of all samples from the virgin yearling heifers (Table 4). From these OTUs, two were classified as *Methanobrevibacter* (OTU8 and OTU23). The remaining shared OTUs were bacterial in origin, with 17% and 80% of them belonging to the *Actinobacteria* and *Firmicutes* phyla, respectively. Of note, three bacterial OTUs [OTU25 (*Eubacterium coprostanoligenes* group; OTU561(*Colidextribacter*) and OTU29 (*Ruminococcus*)] were shared by more than 90% of the samples. In pregnant heifers, 47 OTUs were present in more than 60% of all samples (Table 5), and most of them were identical to those taxa shared among the virgin yearling heifer samples. One taxon identified as *Romboutsia ilealis* (OTU11) was found in 100% of the samples, and OTU24 (*Paeniclostridium*), OTU25, OTU29 and OTU68 (*Bifidobacterium pseudolongum*) were identified in 95% of the samples.

**Table 4.**
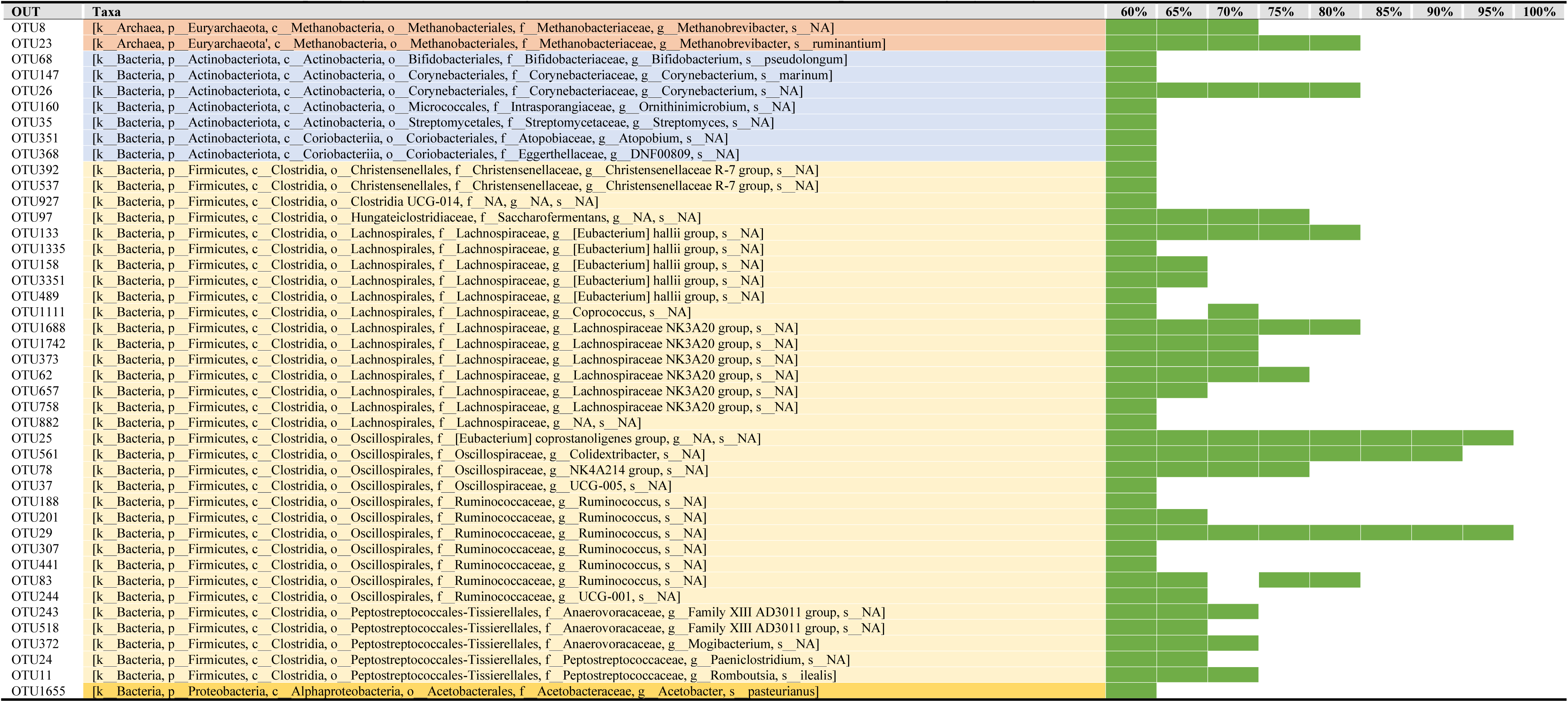
OTUs identified in the nasopharyngeal, ruminal and vaginal microbiota of at least 60% of samples from virgin yearling heifers.

**Table 5.**
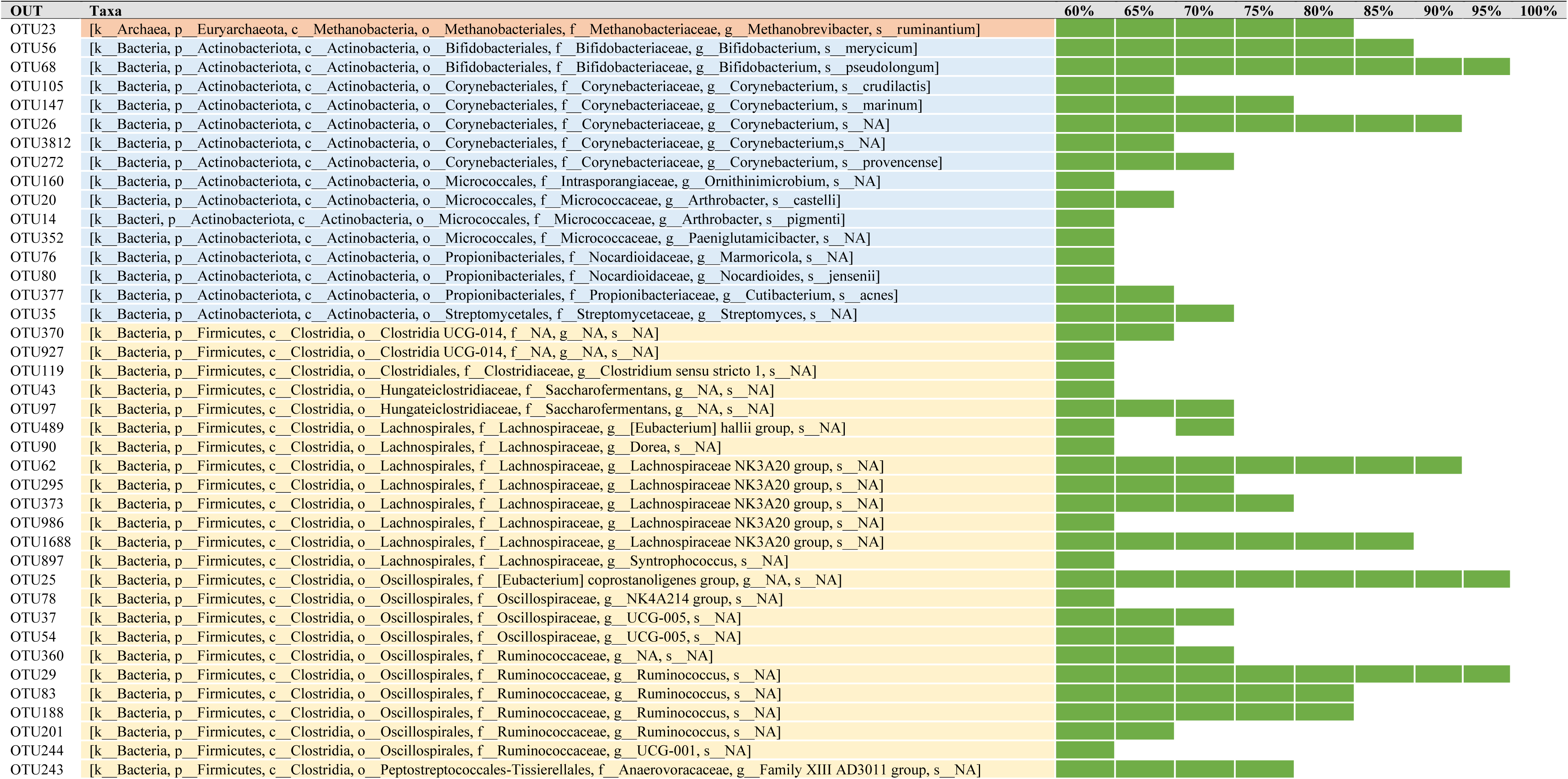
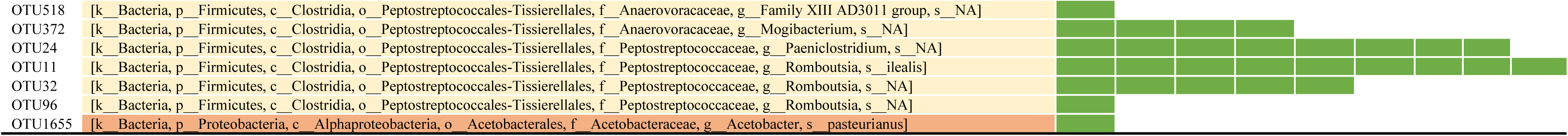
OTUs identified in the nasopharyngeal, ruminal, and vaginal microbiota of at least 60% of samples from pregnant heifers.

**Table 6.**
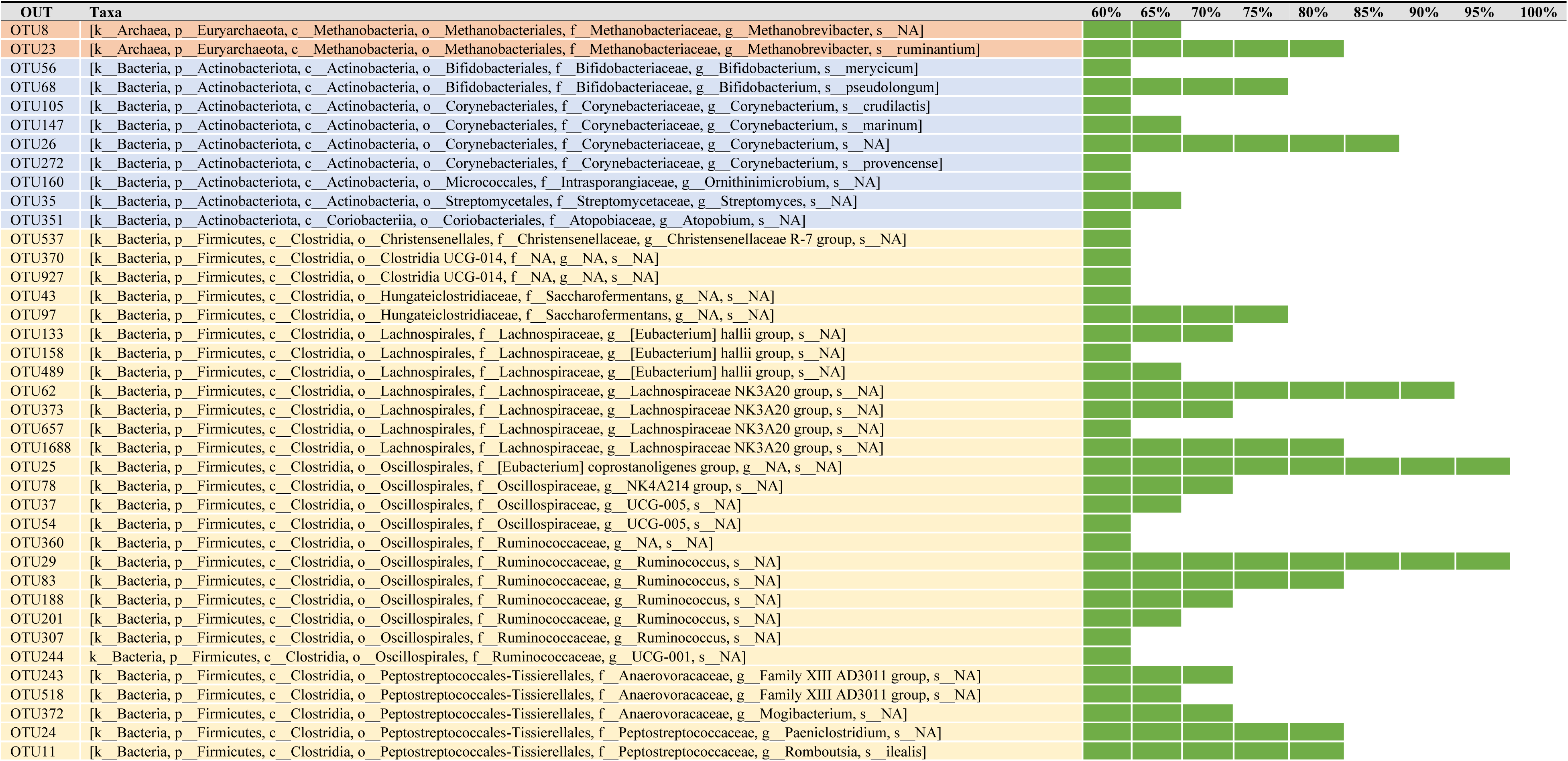
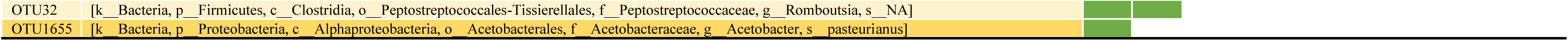
OTUs identified in the nasopharyngeal, ruminal and vaginal microbiota of 60% of samples from yearling and pregnant heifers.

As listed in Table 7, 41 OTUs were identified in 60% of all virgin yearling and pregnant heifer samples. Nine of these OTUs were also present in more than 80% of the samples. These included OTU23 (*Methanobrevibacter ruminantium*), OTU26 (*Corynebacterium*), three OTUs (OTU25, OTU62 and OTU1688) within the *Lachnospiraceae* NK3A20 group, two *Ruminococcus* OTUs (OTU29 and OTU83), OTU24 and OTU11. OTU68 was also found in 75% of the samples. Regardless of animal group and diet, these OTUs were shared by a high proportion of the nasopharyngeal, ruminal fluid, and vaginal samples, suggesting that these taxa may be so-called “core taxa” among these anatomical locations.

### Comparison of Methanogenic Archaeal Relative Abundance Between Virgin Yearling and Pregnant Heifers, and Identification of Bacterial Genera Associated with *Methanobrevibacter*

Methanogenic archaea, and in particular members of the *Methanobrevibacter* genus, have been reported to colonize the intestine of 84-day-old (Amat et al., unpublished), and 5- to 7-month-old calf fetuses (Guzman et al., 2020), as well as newborn calves (Guzman et al., 2015; Alipour et al., 2018). In addition, we identified here two *Methanobrevibacter* OTUs (OTU8 and OTU23) that were shared by a relatively high portion (≥ 65%) of all samples collected from both virgin yearling and pregnant heifers (Table 4). Therefore, we assessed whether the relative abundance of *Methanobrevibacter* changed in response to pregnancy. For this, we compared the overall relative abundance of *Methanobrevibacter* spp. within each sample type between virgin yearling (non-pregnant) and pregnant heifers. Overall, the mean relative abundance of *Methanobrevibacter* in the nasopharynx, rumen fluid and vagina was 0.17%, 5.67%, and 0.47%, respectively (Fig.6A). The relative abundance of *Methanobrevibacter* in the rumen was greater in pregnant heifers compared to yearling heifers (*p* < 0.0001), but similar in the other two microbial habitats (*p* > 0.05) (Fig. 6B and D).

**Figure 6.**
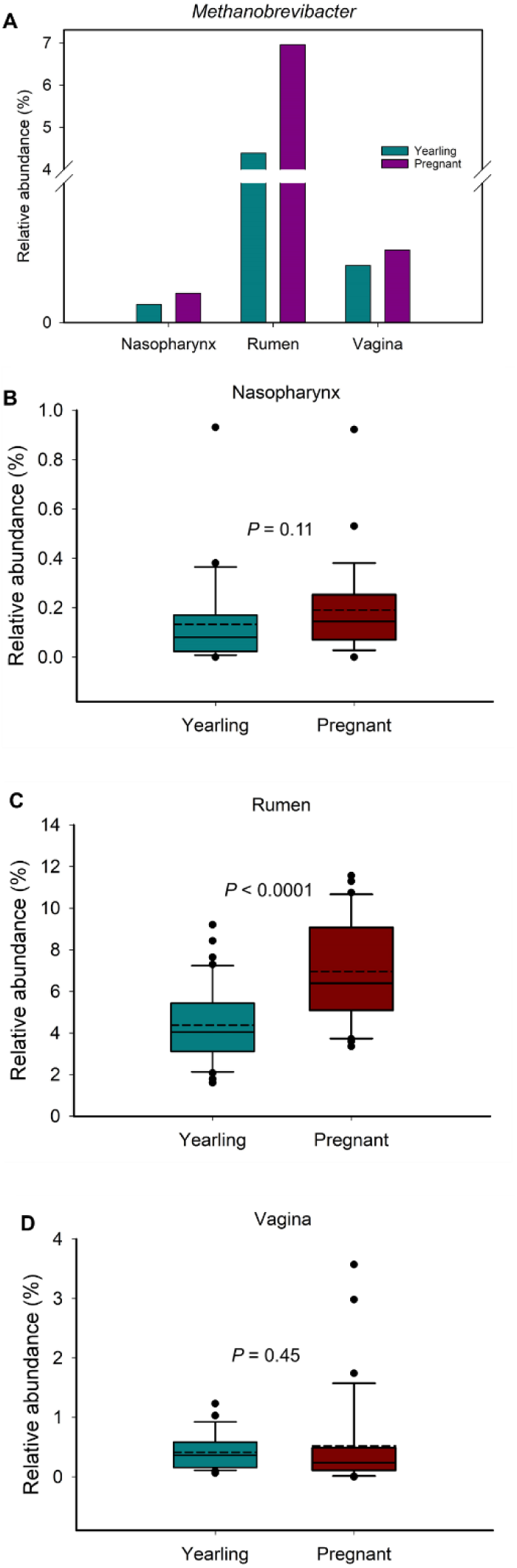
Overall relative abundance of *Methanobrevibacter* in nasopharyngeal, ruminal and vaginal microbiota by sample types (A) and by animal groups (B, C and D).

There is increased research interest in the mitigation of enteric methane emissions from ruminant livestock via the manipulation of the rumen microbiota, primarily targeting commensal bacterial species involved in the supply or consumption of methanogenic substrates. Therefore, we assessed correlations between *Methanobrevibacter* and the other 24 most relatively abundant genera present in vaginal and ruminal microbiota of virgin yearling and pregnant heifers. Of note, the relative abundance of *Methanobrevibacter* in the nasopharynx was relatively low compared to the vagina and rumen and therefore, only genera in the vaginal and ruminal microbiota were included for this correlation analysis. The Spearman correlation analysis revealed that 15 out of these 24 genera in the rumen of virgin yearling heifers exhibited significant (*p* > 0.05) correlations with *Methanobrevibacter*. Among which, the following 10 genera were positively correlated: *Christensenellaceae* R-7 group, *Ruminococcus*, *Oscillospiraceae* NK4A214 group, *Papillibacter*, *Pseudobutyrivibrio*, *Prevotellaceae* NK3B31 group, *Lachnospiraceae* NK3A20 group, *Lachnospiraceae* XPB1014 group, *Eubacterium hallii group*, *Butyrivibrio* and *Olsenella.* Whereas genera within the *Prevotellaceae* family (*Prevotella*, *Prevotellaceae* UCG-003 and *Prevotellaceae* UCG-001) and *Anaeroplasma* were strongly and inversely associated with *Methanobrevibacter* (Fig.7A). Varying degrees of positive or negative associations among the *Methanobrevibacter-*associated 15 genera and between other genera were also identified (Fig.7A). Within the vaginal microbial community of virgin yearling heifers, there were only three genera (*Monoglobus*, *Akkermansia* and *Rikenellaceae* dgA-11 gut group) that displayed significant correlations with *Methanobrevibacter* (*p* < 0.05) and these were positive correlations (Fig. 7B).

**Figure 7.**
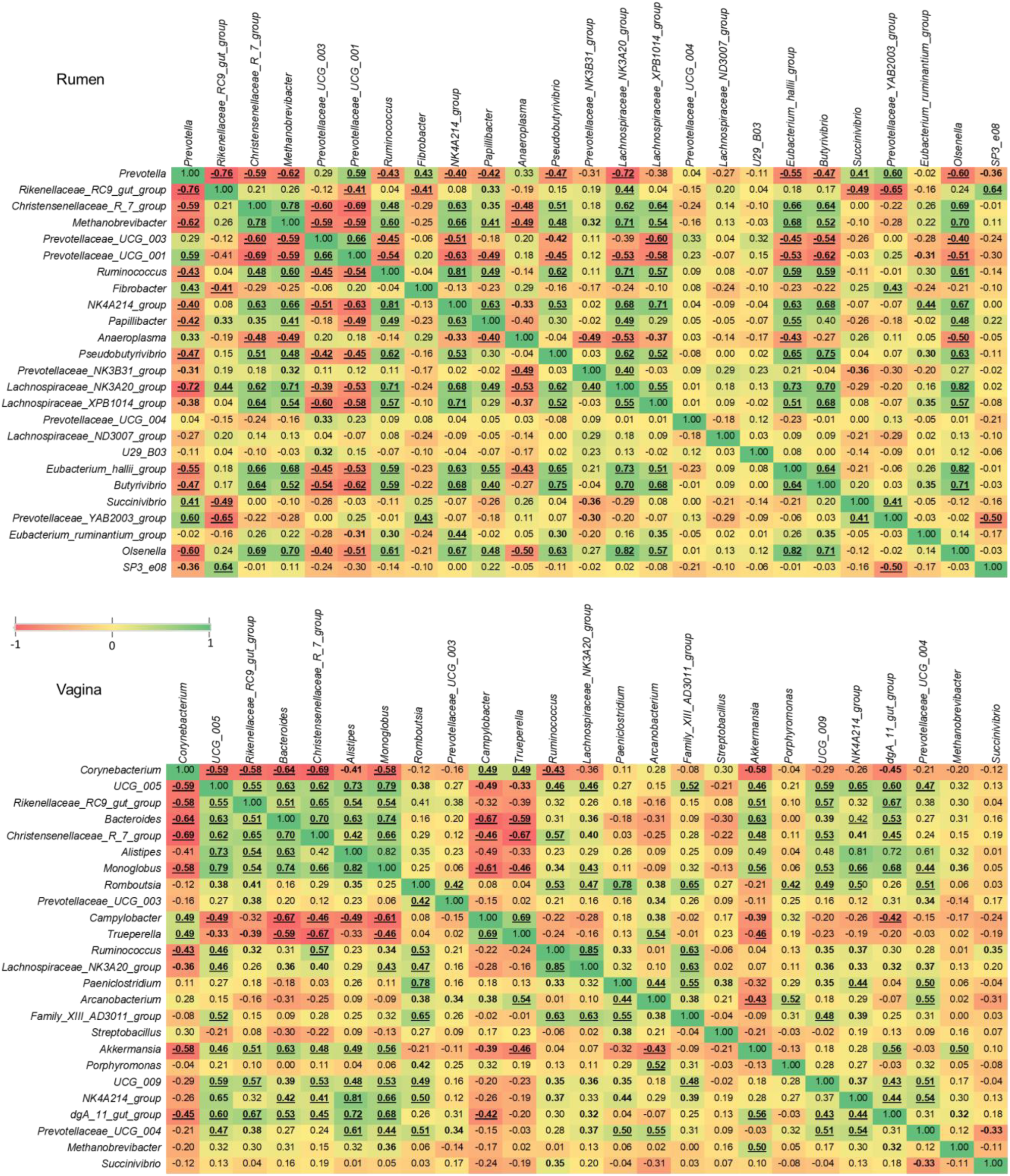
Correlations between the 25 most relatively abundant genera in the ruminal and vaginal microbiota of virgin yearling heifers. Spearman’s rank correlation coefficients (*r*). Bold correlation coefficients with 0.01 ≤ *p* < 0.05, and underlined bold correlation coefficients with *p* < 0.01.

In pregnant heifers, there were similar correlation patterns between *Methanobrevibacter* and other ruminal genera in the yearling heifers, with 14 genera significantly (*P* < 0.05) and positively (n = 8) or negatively (n = 6) correlated with this genus (Fig. 8A). In contrast to the vaginal tract of virgin yearling heifers, there were 10 genera in the vaginal microbiota of pregnant heifers that were significantly associated with *Methanobrevibacter*, nine of them positively correlated. Interestingly, although *Prevotella* and *Prevotellaceae UCG-003* were inversely correlated with *Methanobrevibacter* in the rumen of both virgin yearling and pregnant heifers, they were strongly and positively correlated with this genus in the vagina microbiota. Only the inverse correlations between *Methanobrevibacter* and *Corynebacterium* were significant (*p* < 0.05) (Fig. 8B).

**Figure 8.**
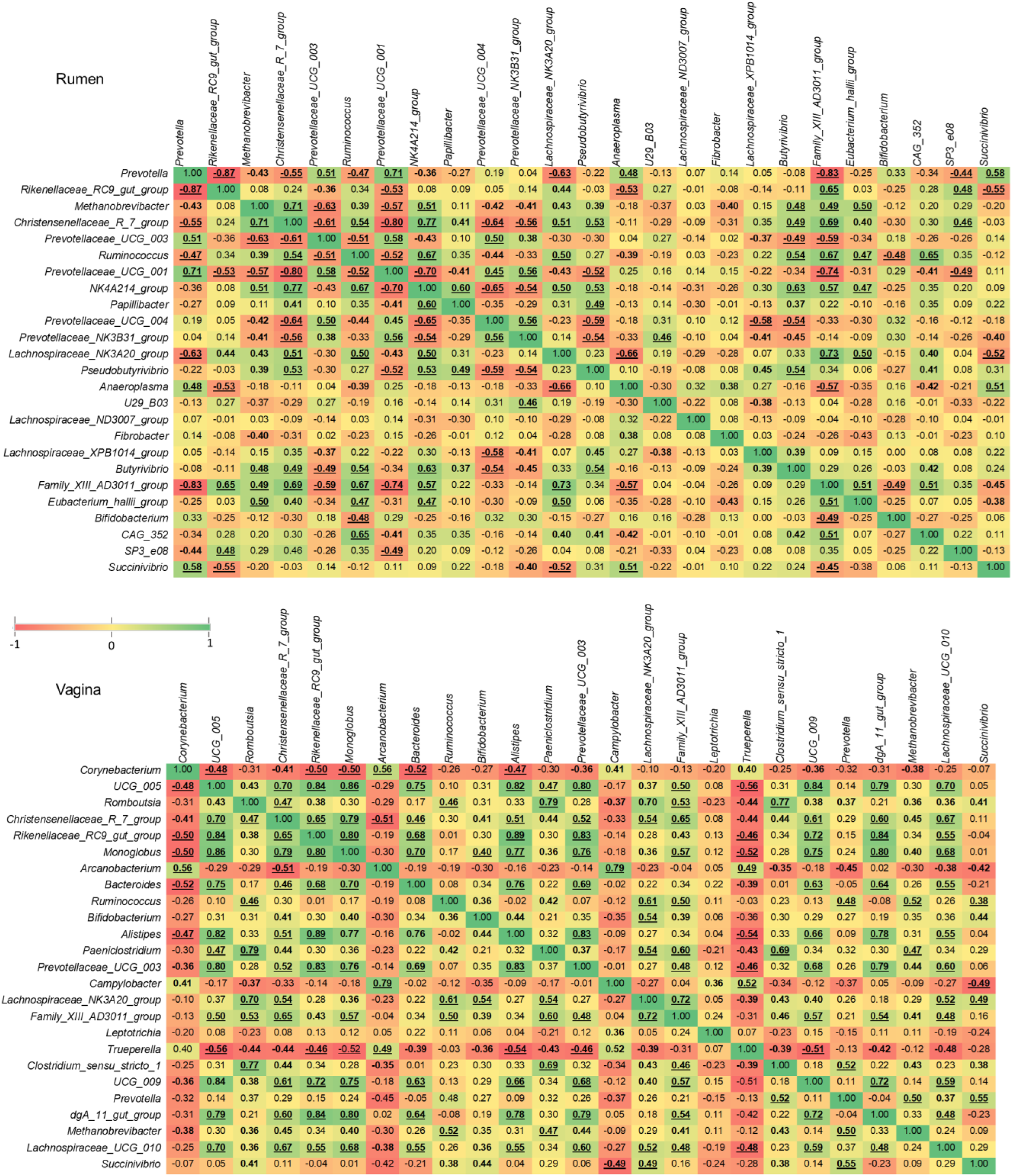
Correlations between the 25 most relatively abundant genera in the ruminal and vaginal microbiota of pregnant heifers. Spearman’s rank correlation coefficients (*r*). Bold correlation coefficients with 0.01 ≤ *p* < 0.05, and underlined bold correlation coefficients with *p*< 0.01.

Next, we applied a stepwise-selected generalized linear mixed model to further identify genera that have a significant effect on the relative abundance of *Methanobrevibacter* spp. in the rumen and vagina. In the virgin yearling heifers, *Prevotella* and the *Christensenellaceae* R-7 group were predicted to have a significant effect (*p* <.0001) on *Methanobrevibacter* [1/(*Methanobrevibacter*^^2^) = 0.02956 + (0.002514 × *Prevotella*) + (-0.00875 × *Christensenellaceae* R-7 group)]. As for pregnant heifers, *Methanobrevibacter* abundance was predicted to be negatively affected by *Prevotellaceae UCG 003* (*p* = 0.037), and positively by *Christensenellaceae* R-7 group (*p* = 0.0326) [1/(*Methanobrevibacter*^^2^) = 0.03793 + (-0.00602 × *Christensenellaceae* R-7 group) + (0.007088 × *Prevotellaceae* UCG-003)]. Within the vaginal microbial community of pregnant heifers, *Ruminococcus* (*p* = 0.0098), *Prevotellaceae* UCG-003 (*p* = 0.0005) and *Prevotella* (*p* <.0001) were predicted to have a significant negative effect on *Methanobrevibacter* [1/(*Methanobrevibacter*^^2^) = -0.3780 + (0.1501 × *Ruminococcus*) + (0.2875 × *Prevotella* UCG-003) + (0.2898 × *Prevotella*)]. Among the 24 most relatively abundant genera in the vaginal microbiota of yearling heifers, only *Oscillospiraceae* UCG-005 (*p* = 0.0138) was predicted to have a negative impact on *Methanobrevibacter* [1/(*Methanobrevibacter*^^2^) = 0.05975 + (0.04037× *Oscillospiraceae* UCG-005).

## DISCUSSION

New evidence from our laboratory (Amat et al., unpublished data) and Guzman and colleagues (Guzman et al., 2020) indicate that microbial colonization of the calf gastrointestinal tract may take place before birth. These observations suggest that the maternal microbiome may have a role in shaping the development of the offspring microbiome in cattle. In addition, it is believed that undesirable alterations of the maternal microbiota may indirectly influence fetal development, and that these effects may be transmitted to progeny, resulting in a dysbiotic microbiota (Calatayud et al., 2019) and increased offspring susceptibility to the development of metabolic disorders and respiratory infections (Calatayud et al., 2019; Yao et al., 2020). Recent evidence from mouse studies has demonstrated that the maternal microbiota during pregnancy modulates the programming of fetal metabolic and nervous system development (Kimura et al., 2020; Vuong et al., 2020).

Although the role of maternal nutrition in developmental programming in cattle has been relatively well appreciated (McLean et al., 2017; Caton et al., 2019; Crouse et al., 2019; Menezes et al., 2021), the potential involvement of the maternal microbiota in fetal programming and offspring microbiome development remains largely unexplored. Considering the current evidence, it is important to explore whether bovine maternal nutrition/microbiome during pregnancy influences feto-maternal crosstalk, subsequently influencing offspring microbiome development. Maternal vitamin and mineral supplementation before calving has been well documented to be associated with improved fetal programming and offspring health and productivity in cattle (Mee et al., 1995; Wilde, 2006; Van Emon et al., 2020; Diniz et al., 2021; Menezes et al., 2021). However, whether VTM supplementation-associated positive outcomes observed pre- and post-calving are dependent on alterations in the ruminal microbiota remains unexplored. In the present study, we evaluated whether differences in maternal weight gain during the first trimester of gestation affected the postnatal nasopharyngeal, ruminal, and vagina microbial communities of virgin heifers at 9 months of age. We also characterized and compared these three microbiota in pregnant heifers to evaluate the impact of VTM supplementation during the first six month of gestation on the maternal microbiome. Finally, we identified core taxa that are shared within the respiratory, gastrointestinal, and reproductive tract microbiota of cattle.

### Effect of Maternal Restricted Gain During the First Trimester of Gestation on Offspring Microbiota Development

The virgin yearling heifers born from the dams that were subjected to LG (0.29 kg/d) during the first 84 days of gestation harbored a similar nasopharyngeal, ruminal and vaginal microbiota to those born from MG (0.79 kg/d) dams. The LG dams had a reduced average daily gain (*p* < 0.01) were 40 kg lighter than MG dams at calving (*p* < 0.01), and their calves had a lower birth weight than those from MG dams (28.6 vs. 30.8 kg, *p* = 0.03) (Baumgaertner, 2020). As previously reported, fetuses harvested from a subset of the LG and MG dams at 84 days of gestation exhibited distinct fetal metabolic programming (Menezes et al., 2021), including altered amino acid profiles in the fetal fluids (Menezes et al., 2021). In addition, we identified the presence of an archaeal and bacterial microbiota in intestinal and fluid samples from these 84-day-old calf fetuses (Amat et al., unpublished data). Therefore, we hypothesized that the divergent microbiome may be detected in virgin heifers that were exposed to divergent in utero nutrition (i.e. LG or MG) during their first trimester of gestation. However, no significant effect of maternal nutrition was found on the microbial community structure in the offspring nasopharynx, rumen, or vagina. There may be many reasons for this finding, including the timing of sample collection. For example, samples were collected when the offspring heifers were at 9 months of age, which was about 15 months after fetal exposure to the restrictive maternal diets. This may simply be too late in their development to detect microbial community alterations in the offspring as a result of maternal nutrition. Therefore, future studies investigating the impact of maternal nutrient and microbiome on offspring microbiome development should include a more robust profile of early life microbiome measurements.

### Effect of Vitamin and Mineral Supplementation During the First 6 Months of Gestation on Maternal Microbiota

VTM supplementation during the first 6 months of gestation did not induce significant alterations in community structure and diversity of the nasopharyngeal, rumen or vaginal microbiota. While there is limited information on the effect of mineral and vitamin supplementation on the gut microbiota of ruminant animals, the impact of dietary mineral and vitamin intake on potentially beneficial or pathogenic gut microbes in humans and rodent animals have been relatively well documented (Yang et al., 2020). For example, calcium and phosphorus supplementation increased the relative abundance of *Clostridium*, *Ruminococcus* and *Lactobacillus* spp. while reducing *Bifidobacterium* spp. in healthy men or mice (Nadeem Aslam et al., 2016; Trautvetter et al., 2018; Li et al., 2019). The impact of dietary supplementation with selenium, magnesium, iron or zinc on certain gut commensal or pathogenic microbes was also reported in children and mice (Yang et al., 2020). Of note, a significant effect of mineral supplementation on the gut microbiota was observed but only at the microbial taxa level and not on the microbial community structure and diversity (Yang et al., 2020).

The results from a limited number of studies performed on cattle also indicate that mineral supplementation may influence ruminal microbiota composition. Clay mineral supplementation increased the relative abundance of *Butyrivibrio* while reducing the relative abundance of *Lactobacillus*, *Fusobacterium,* and *Treponema* genera in the rumen of non-lactating Holstein cows (Neubauer et al., 2019); however, it did not alter the rumen microbial community structure or diversity. Similarly, we observed that VTM supplementation increased the relative abundance of *Butyrivibrio* in the rumen (*p* < 0.05). *Butyrivibrio* spp. are considered commensal members of the rumen microbiota, producing butyrate through degradation of otherwise indigestible plant polysaccharides (Kelly et al., 2010). The *Oscillospiraceae* NK4A214 group and *Succinivibrionaceae* CAG-352 were the only other ruminal genera that responded to VTM supplementation in the present study. These are uncultured taxa and their role in the rumen is largely unknown. In contrast to our findings and those of Neubauer et al. (2019), Liu and others (2017) observed noticeable alterations in microbial richness and diversity of ruminal microbiota in both lactating Holstein cows (3-4 years old) and yearling heifers (10-months old) in response to feeding mineral salt bricks containing Mg, Co, Cu, Fe, Mn, Se, Zn, I and Na for one month.

Compared to the rumen microbiota, the effect of mineral supplementation on the bovine respiratory and reproductive microbiota ha been less characterized. Feeding beef calves with selenium-biofortified alfalfa hay has been reported to alter the nasopharyngeal microbiota (Hall et al., 2017; Hall et al., 2020). In the present study, although no significant changes were detected in the microbial community structure and diversity in the nasopharynx following VTM supplementation, changes were detected in relative abundance of five relatively abundant genera (*Mycoplasma*, *Oscillospiraceae* UCG-005, *Christensenellaceae* R-7 group, *Ruminococcus* and *Ornithinimicrobium*). Among these genera, *Mycoplasma*, which includes a bovine respiratory disease (BRD)-associated pathogen, *Mycoplasma bovis*, was enriched in pregnant heifers that receiving VTM supplementation. BRD is not a significant health concern among adult and pregnant cattle but it is the number one health problem affecting newly weaning calves arriving in the feedlot (Johnson and Pendell, 2017). The positive association between VTM supplementation and nasopharyngeal *Mycoplasma* observed here poses the question of whether maternal VTM supplementation influences colonization of the offspring respiratory tract by *Mycoplasma* spp. No information has been reported regarding the impact of mineral supplementation on reproductive microbiota in cattle.

Vitamins A, D_3_ and E were included in the VTM supplement given to the pregnant heifers. Thus, it is impossible to discern whether the subtle changes observed at the taxa level in both the nasopharyngeal and ruminal microbiota are due to the minerals and vitamins supplemented. Evidence from human, rodent and pig studies suggest that the gut microbiota responds to vitamin supplementation (Yang et al., 2020). Gastrointestinal-associated *Bifidobacterium* (vitamin A, C) *Akkermansia* (vitamin A), and *Lactobacillus* spp. (vitamin C) were more relatively abundant while *E. coli* (vitamin C) and *Clostridium* (vitamin D) spp. decreased in relative abundance after vitamin supplementation (Xu et al., 2014; Talsness et al., 2017; Huda et al., 2019; Yang et al., 2020). Our results indicate that vitamin supplementation has less impact on the ruminal microbiota. Overall, VTM supplementation for first 6 months of gestation did not affect the maternal microbiota. There could be due to several factors. Considering that mineral salt intake was reported to alter the ruminal microbiota in 3-4-year-old lactating cows (Liu et al., 2017), the resilience and robust of the mature ruminal microbiota can likely be ruled out as a reason for the absence of any VTM effect on the ruminal microbiota in pregnant heifers (1 year 9 months-old).

Pregnancy status rather than age, however, could be associated with the non-responsiveness of the ruminal microbiota to VTM supplementation. In rodent studies, the maternal gut microbiota undergoes profound changes over the course of pregnancy (Collado et al., 2008; Koren et al., 2012; Nuriel-Ohayon et al., 2016; Smid et al., 2018). As pregnancy progresses from the 1st to 3rd trimester, the maternal gut microbiota becomes less diverse (Koren et al., 2012) but with a higher microbial density, which may result in a microbiota that is more robust and resilient to perturbations. Hence, future studies are warranted to investigate the impact of VTM supplementation and other dietary interventions on the maternal microbiota of cattle using a non-pregnant control cohort.

### Holistic View of Microbial Communities Across Respiratory, Gastrointestinal and Reproductive Tract and the Core Taxa Shared Across These Habitats

As expected, the overall microbial structure, diversity and composition were noticeably different among the nasopharyngeal, ruminal and vaginal microbiota in both virgin yearling and pregnant heifers. The ruminal microbiota was dominated by the anaerobic phylum *Bacteroidota* (66%), while the nasopharyngeal and vaginal microbiota the majority of 16S rRNA gene sequences were classified as *Actinobacteriota* (51%) and *Firmicutes* (52%), respectively. Various factors including niche-specific physiological factors (temperature, pH, oxygen and nutrient availability), dietary, and environmental factors are involved in shaping the microbiota of the bovine respiratory tract (Zeineldin et al., 2019; Timsit et al., 2020), rumen (O’Hara et al., 2020; Cholewińska et al., 2021) and reproductive tract (Galvão et al., 2019). Subtle physiological and anatomical differences in the mucosal surfaces of the bovine respiratory tract have been shown to significantly influence the microbial distribution along the respiratory tract (McMullen et al., 2020).

In the present study, although the nasopharynx, rumen and vagina have drastically different physiological and anatomical properties, we identified 41 OTUs that were shared by a high portion (60%) of all samples from both virgin yearling and pregnant heifers. This indicates that these core taxa can colonize and inhabit the respiratory, gastrointestinal and reproductive tracts regardless of the drastic differences in physiological conditions in these locations. The majority (80%) of these core taxa are members of the *Firmicutes*, which is one of the most ubiquitous and relatively abundant bacterial phyla in the respiratory, gastrointestinal and reproductive tract- (vagina, uterus) (Galvão et al., 2019), mammary gland- (Derakhshani et al., 2018), ocular- (Bartenslager et al., 2021) and hoof- (Zinicola et al., 2015) -associated microbiota in cattle, demonstrating the adaptability of members of this phylum. Nine taxa within the *Actinobacteria* phylum including *Bifidobacterium pseudolongum* and several *Corynebacterium* spp. were among these core taxa. *B. pseudolongum* is widely found in the mammalian gut (Lugli et al., 2019) and has long been noted for its probiotic properties in human, cattle and pigs (Abe et al., 1995; Kissels et al., 2017). Given the known beneficial effects of this species on the host, and as a core taxon present in the respiratory, gut and reproductive tracts of cattle, *B. pseudolongum* may have the potential to enhance cattle health and productivity, as may some of the other core taxa identified in this study. Species and strain level resolution of these core taxa using shotgun metagenomic sequencing and characterization of their functional features using culturing should be the focus of future studies.

Two *Methanobrevibacter* OTUs were also identified among the core taxa. Although members of this methanogenic genus are well known for their involvement in ruminal methane production (Hook et al., 2010; Danielsson et al., 2017; Greening et al., 2019), and are frequently observed in the vaginal microbiota (Laguardia-Nascimento et al., 2015) in cattle, it is interesting to note that this genus is also found in the bovine respiratory tract. The presence of these *Methanobrevibacter* OTUs within the respiratory, gastrointestinal, and reproductive tracts has important implications for the identification of maternal seeding of the calf microbiota with pioneer methanogens before and during birth. *Methanobrevibacter* spp. are predominant in 5-to 7-month-old calf fetuses (Guzman et al., 2020) as well as newborn calves (Guzman et al., 2015; Alipour et al., 2018). Our results indicate that the respiratory microbiota may also seed the calf gastrointestinal tract with *Methanobrevibacter* spp. perinatally. This highlights the necessity of holistic assessment of respiratory, gastrointestinal and reproductive tract microbiota to trace the origin of pioneer calf microbiota. To our best of knowledge, this is the first study to evaluate nasopharyngeal, ruminal and vaginal microbiota in an individual ruminant animal, and to identify the core taxa shared amongst these microbial ecologies.

### Ruminal *Methanobrevibacter* Enriched in Pregnant Heifers and Associations of *Methanobrevibacter* with Predominant Ruminal and Vaginal Bacterial Genera

Given that lowering methane emissions in cattle benefits both environment and cattle production (Beauchemin et al., 2020), and increasing evidence suggesting that the ruminal microbiome and host genetics can be targeted independently to improve feed efficiency and mitigate enteric methane emissions from cattle (Difford et al., 2018; Li et al., 2019), we therefore focused on this methanogenic archaeal genus, *Methanobrevibacter*. We identified that pregnant heifers harbored a greater relative abundance of ruminal *Methanobrevibacter* compared to non-pregnant virgin yearling heifers. Confounding factors associated with dietary (11 % more hay fed to virgin heifers than pregnant heifers) and age differences makes it difficult to associate pregnancy with the colonization of the rumen with methanogenic archaea. However, the impact of pregnancy and mitigation of maternal ruminal methanogens on offspring enteric methane emissions warrants further investigation.

Our correlation analysis revealed that in comparison to vaginal *Methanobrevibacter*, the relative abundance of ruminal *Methanobrevibacter* is highly influenced by many other commensal genera in the rumen. For example, in the rumen microbiota many genera within the *Prevotellaceae* family were inversely associated with the relative abundance of *Methanobrevibacter.* Interestingly, the opposite was found in the vaginal microbiota, suggesting that the nature of the interaction between *Methanobrevibacter* and *Prevotella* and *Prevotellaceae* UCG-003 may be niche specific and that *Prevotella* spp. in the rumen may become pro-methanogenic if they present in reproductive microbial ecosystem.

The stepwise-selected GLM model identified *Prevotella* and *Prevotellaceae UCG-003* as having a significant and negative effect on the relative abundance of *Methanobrevibacter* in rumen of both virgin and pregnant heifers and vaginal tract of pregnant heifers. This is in agreement with previous studies reporting that microbial communities with highly-abundant lactate-consuming bacteria (*Prevotella bryantii*) and high H_2_-consuming (e.g. certain *Prevotella* spp.) has been associated with lower ruminal methane production (Denman et al., 2015; Danielsson et al., 2017; Tapio et al., 2017; Granja-Salcedo et al., 2019). Thus, members of the *Prevotella* and *Prevotellaceae UCG-003* in the bovine rumen and vagina may have anti-methanogenic potential to mitigate methane emissions in cattle. The *Christensenellaceae R-*7 group was identified as the genus that can have significant positive effect on *Methanobrevibacter* in the present study. This may suggest that some species within this genus may be involved in producing methanogenic substrates such as H_2_ and acetate. Future *in vitro* studies are needed to confirm the anti-methanogenic properties of *Prevotella* and *Prevotellaceae UCG-003* and pro-methanogenic activity of *Christensenellaceae R-7* group spp. originating from the rumen of cattle.

In conclusion, no noticeable difference was observed in α and β-diversity in any of the nasopharyngeal, ruminal and vaginal microbiota between virgin heifers raised from dams exposed to divergent rates of gain during the first trimester of pregnancy, or between pregnant heifers consuming control and VTM diets. Only in the vaginal microbiota were there relatively abundant genera that were affected by maternal rate of gain during early gestation. Maternal VTM supplementation resulted in subtle compositional alterations in the nasopharyngeal and ruminal microbiota. A total of 41 archaeal and bacterial OTUs were shared by over 60% of all samples from both virgin and pregnant heifers. Two taxa within the *Methanobrevibacter* genus were among these taxa this genus was more relatively abundant in pregnant compared to virgin heifers. Compared to the vaginal *Methanobrevibacter*, *Methanobrevibacter* in the rumen was predicted to be highly interactive with other commensal members.

Among the 25 most relatively abundant genera, *Prevotella* and *Prevotella* UCG-003 (negative) and *Christensenellaceae R-7* group (positive) were predicted to have a significant effect on the relative abundance of ruminal *Methanobrevibacter* spp. Overall, the results of this study suggest that there is little impact of maternal gestational nutrition during the first trimester on the calf microbiota assessed at 9 months of age, and that VTM supplementation during pregnancy may not alter the maternal microbiota. This study provides evidence that there are several microbial taxa, including methanogenic archaea, that are shared across the respiratory, gastrointestinal, and reproductive tracts. Therefore, this suggests that there is a need for a holistic evaluation of the bovine microbiota when considering potential maternal sources for seeding calves with pioneer microbiota, and when targeting the maternal microbiome to enhance offspring health and development.

## DATA AVAILABILITY

The datasets generated for this study can be found in the Sequences that were submitted to the NCBI sequence read archive under BioProject accession PRJNA721423.

## ETHICS STATEMENT

All animals used in this study were cared for in accordance with the guidelines set by the Olfert et al. (1993) and all experimental procedures involving cattle were approved by the North Dakota State University Institutional Animal Care and Use Committee (#A20085 and A20047).

## AUTHOR CONTRIBUTIONS

Conceiving the idea, designing the study, and providing supervision—S.A. and C.R.D.; Cattle management— A.C.B.M., F.B., C.R.D. and K.K.S.; Animal care and sample collections— S.A., C.R.D. K.S., A.C.B.M., T.W., J. D. K., F.B.; Sample processing—K.S. and S.A.; Bioinformatics analysis— D.B.H., T.L. and S.A.; Data processing and statistical analysis—T.D.S., D.B.H, and S.A.; Manuscript writing—S.A.; Manuscript review and editing—S.A., D.B.H., A.C.B.M and C.R.D. All authors have read and agreed to the published version of the manuscript.

## ACKNOWLEDGMENTS

The authors acknowledge the support from the staff at NDSU Beef Cattle Research Complex.

